# The Ocean Microbiomics Database reveals thermal niches and projected pressure on marine microbes in a warming ocean

**DOI:** 10.64898/2026.09.28.755000

**Authors:** Samuel Miravet-Verde, Hans-Joachim Ruscheweyh, Kang Li, Simona Skiotytė, Taylor Priest, Anna Sintsova, Luca Schnepp-Pesch, James O’Brien, Harun Mustafa, André Kahles, Lucas Paoli, Georg Zeller, A. Murat Eren, Shinichi Sunagawa

## Abstract

Temperature is a key driver of ocean microbial community structure, yet its role in shaping global distributions and their response to warming remains unclear due to fragmented studies, inconsistent methods, and limited environmental context. We present the Ocean Microbiomics Database (OMDB) containing 274,282 genomes forming 31,058 species-level clusters, searchable at https://omdb.microbiomics.io. We show surface temperature is predicted from community composition, driven by thermal niche partitioning among congeneric taxa. Using these thermal niches to model present-day distributions globally, we find that projected warming results in microbial richness declining faster than relative abundance, with losses concentrated at high latitudes and among poorly characterized lineages. Resolving 5,891 species into four thermal clusters reveals ecological structure correlated with genome streamlining, protein thermostability, and marker-gene physicochemistry, enabling prediction of a microorganism’s abundance-maximizing temperature directly from its coding sequence. This positions OMDB as a platform spanning molecular traits to global ecology, offering a genome-resolved foundation for understanding how temperature, shown to be a primary driver of ocean microbial life, will continue to shape it as the ocean warms.

## Introduction

Ocean warming has been linked to shifts in microbial community composition^1,2^, function^3^ and diversity^4^, as well as reduced primary productivity^5^. Driven primarily by human activities, the global ocean surface has already warmed by 0.88 °C relative to the pre-industrial baseline (very likely range: 0.68 to 1.01 °C)^6^. However, whether warming leads to compensatory shifts toward more resilient communities or the loss of key taxa^7,8^ remains unclear. More broadly, growing recognition of the vulnerability of microorganisms to global change has prompted new initiatives to safeguard microbial biodiversity^9^.

The biodiversity at stake is immense: the global ocean microbiome encompasses extensive phylogenetic and functional diversity, and underpins biogeochemical cycling, ecosystem functioning, and climate regulation^1,10,11^. However, studying these communities remains challenging due to their complexity and the difficulty of cultivating most microorganisms under laboratory conditions^12^. Culture-independent approaches have therefore become indispensable for accessing and analyzing the relationships between microbial community structure and function^13^. More recently, advances in gene-centric and genome-resolved approaches have enabled the discovery of previously unknown microbial taxa, enzymes and bioactive molecules with potential applications in biotechnology, biomedicine, and synthetic biology^14,15^. Yet each of these discoveries also underscores that we are only beginning to grasp the extent of microbial diversity and genetic resources in the global ocean, much of which may be reshaped, or lost, under ongoing environmental change before it has been described.

Assessing how environmental change will affect this diversity first requires understanding which environmental conditions shape microbial communities, and temperature is a prime candidate, particularly in the photic zone, where primary production sustains marine food webs^16^. Although temperature has been proposed as a key driver of ocean microbial community structure^1^, the molecular basis of this relationship, its robustness across studies, and its global generalizability remain unresolved. For example, classical indicators of thermal adaptation, such as amino acid frequencies^17^ or GC content^18^, do not consistently generalize across marine systems and remain difficult to interpret mechanistically. It also remains unclear whether this structuring is reflected in the partitioning of individual microbial groups into distinct thermal niches, and whether such niches could meaningfully contextualize microorganisms within the emerging concept of climate-impacted changes in microbial biodiversity^19^. Although physiological and genomic correlates of thermal adaptation have been described for individual taxa^7,8,20,21^, addressing these questions requires scaling temperature-response analyses to an ocean microbiome-wide framework. Such a framework should ideally link ecological patterns directly to the proteomic and structural variations that govern thermal tolerance^22^, and extend insights from individual taxa and localized studies to global comparisons that maximize taxonomic range and environmental gradients.

Implementing such a framework, however, is limited by the current state of genomic resources. Several data collections comprise isolate, single-cell, or metagenome-assembled genomes (MAGs) with inconsistent contextual information, hindering comparative analyses. Differences in assembly algorithms (*e.g.*, metaSPAdes^23^ vs. MEGAHIT^24^), reconstruction strategies (*e.g.*, single-sample- vs. co-assembly, and the use of contig coverage only within or across samples^25,26^), and parameter settings can substantially affect the quantity and quality of recovered genomes^27^, further complicating integrative analyses across studies. Comparability is further limited by data fragmentation across repositories. General resources, such as MGnify^28^ or GlobDB^29^, aggregate MAGs produced using heterogeneous methods, whereas ocean-specific databases like OceanDNA^30^ and GOMC^15^ offer limited support for taxonomy, geography or sequence-based queries and lack links to original data sources. Consequently, tracing genomes back to their samples of origin, associated raw data, and contextual metadata remains challenging, leaving critical information scattered across databases and publications. Furthermore, genome collections alone do not provide metagenomic abundance profiles that are needed to study ecological distributions, biogeographic patterns, environmental associations, and population dynamics^31^. Finally, the lack of a unified platform for genome, abundance, and sequence searches limits the ability to place new ocean microbiome data in a global context, to assess their novelty and distributions in light of a changing ocean.

Here, we present the Ocean Microbiomics Database (OMDB), a genome-resolved resource of integrated ocean metagenomes across the globe, giving access to microbial genomes by taxonomy, geography and sequence at https://omdb.microbiomics.io. We analyzed over 270,000 microbial genomes recovered from 12,000 metagenomes and show that sea surface temperature can be accurately predicted from microbial composition profiles independent of methodological biases and geographic proximity, revealing closely related but ecologically diversified taxa among the main predictors. By aggregating temperature profiles of all ocean microbiome members, we further reveal patterns of thermal niche partitioning and geographically map microbial groups that are projected to encounter temperatures beyond their present-day realized niches under ocean warming. Finally, we show that thermal preferences can be linked to genome streamlining and protein thermostability and predicted from physicochemical protein descriptors. Together, our work demonstrates how OMDB facilitates the study of ocean microbial life across scales, from molecular traits to global ecological patterns, and establishes a baseline to contextualize the ocean microbiome within a framework of climate-threatened microbial biodiversity.

## Results

### OMDB enables interactive and integrated exploration of marine microbial genomes through taxonomy, geography and sequence searches

To address the fragmentation, inconsistent processing, and sparse annotation of ocean microbiome data, we build on previous work^14^ to establish the OMDB. This resource substantially expands existing resources through the uniform reconstruction of genomes from globally distributed sampling sites and depths, and provides an interactive web interface for exploring the data by taxonomy, geography, and sequence.

We first selected 209 ocean microbiome studies, including major campaigns such as *Tara* Oceans^32^, *Tara* Pacific^33^, Bio-GO-SHIP^34^, bioGEOTRACES, the Bermuda-Atlantic Time-series Study, and the Hawaii Ocean Time-series^35^, resulting in a comprehensive geographic coverage of the global ocean (Figure 1A). We then downloaded the associated metagenomic data (n = 12,347) from the European Nucleotide Archive (ENA) and metadata from both ENA records and corresponding publications. Specifically, we manually curated key sample metadata (geolocation, depth, collection date, size fraction, and temperature) by cross-checking source publications and primary repositories to standardize data units and formats. We also extracted Copernicus Marine Service environmental variables (salinity, oxygen, nitrate, phosphate, pH, and phytoplankton biomass)^36^ based on sample coordinates and collection dates, yielding a comprehensive, quality-controlled metadata table (Figure S1).

**Figure 1.**
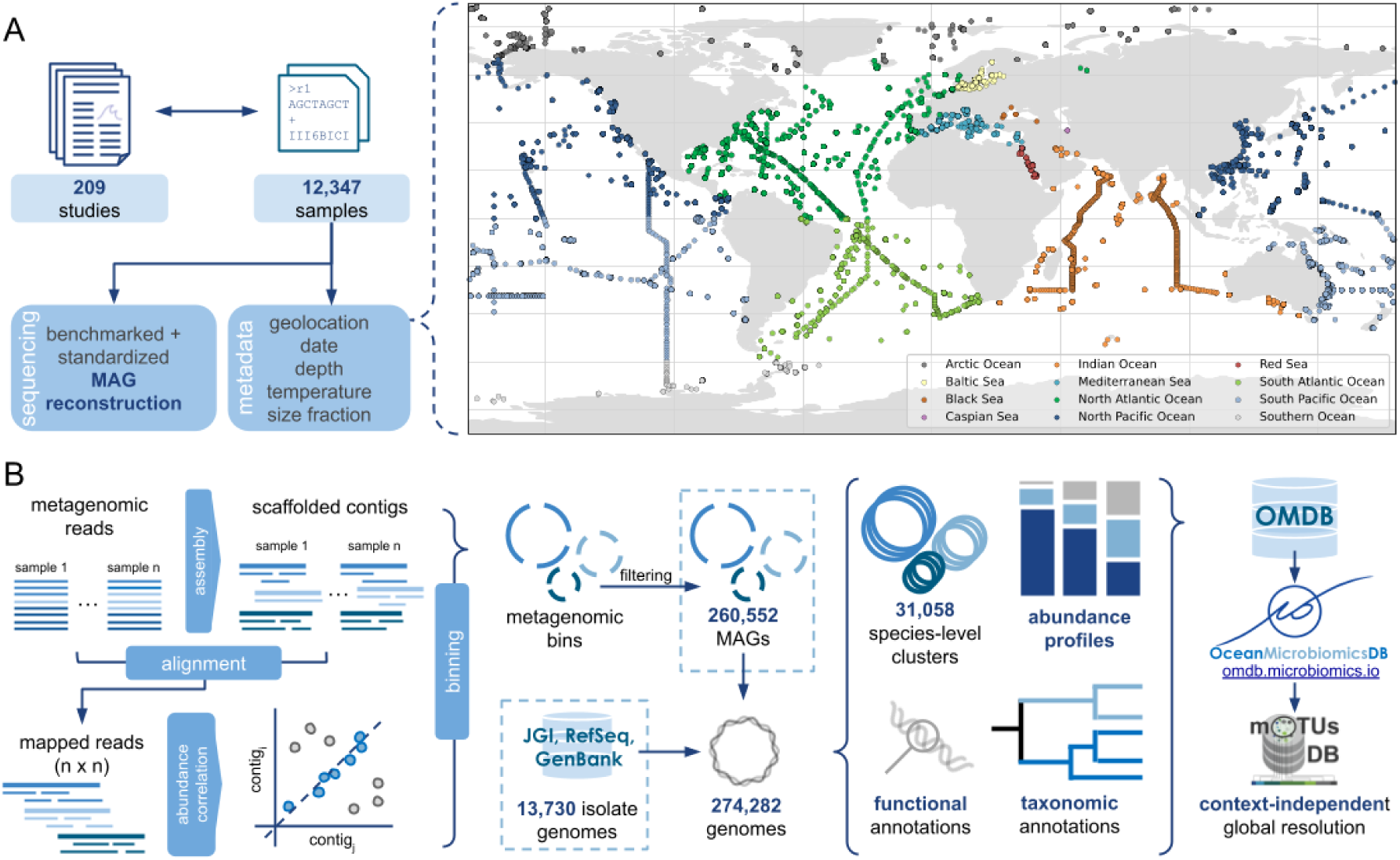
Standardized genome reconstruction and global representation of the Ocean Microbiomics Database (OMDB). (**A**) Overview of sample and metadata curation. Left: metagenomic samples from ocean microbiome studies were retrieved from public repositories and matched with metadata to enable standardized and benchmarked metagenome-assembled genome (MAG) reconstruction. Right: global distribution of sampling sites included in OMDB, colored by ocean/sea region (see inset). (**B**) Genome reconstruction and OMDB integration workflow. Quality-controlled metagenomic reads from each sample were assembled into scaffolded contigs and binned into MAGs using quality-enhancing cross-sample contig coverage correlations. These were combined with isolate and single-cell amplified genomes from JGI, RefSeq, and GenBank to produce a final set of 274,282 ocean microbial genomes. These genomes were clustered into species-level clusters, taxonomically and functionally annotated, and integrated into the OMDB web platform (https://omdb.microbiomics.io). OMDB is designed to be fully compatible with the mOTUs online database (https://motus-db.org)^39^, providing cross-environmental context for ocean microbiome data.

To improve binning performance and thereby enhance genome recovery, we used a standardized and previously benchmarked workflow that groups contigs in individual samples into metagenome assembled genomes (MAGs) by leveraging the co-variance of contig abundances across samples^14,37^. In total, we processed 95.7 TB of compressed sequencing data, encompassing approximately 1.33 trillion reads and 173 trillion nucleotides to reconstruct 260,552 MAGs. Among these, 57,967 OMDB MAGs were labelled as high-quality (≥90% completeness and ≤5% contamination) and 91,706 as good-quality (≥70% completeness and ≤10% contamination) (Figure S2). To further enrich the coverage of genomic information, 13,730 publicly available marine isolate and single-cell amplified genomes from RefSeq, GenBank and JGI portal^38^ were incorporated, yielding a final dataset of 274,282 ocean microbial genomes (Figure 1B). Together, OMDB greatly exceeds previous integrative attempts for a genomic representation of the global ocean microbiome (Figure S2), and is fully integrated into a global genomic data framework^39^ to facilitate cross-ecosystemic contextualization of the ocean microbiome (Figure S3). Furthermore, for each genome, we applied state-of-the-art bioinformatic pipelines to predict genes, functional RNAs, and proteins, alongside both general (KEGG, PFAM, EggNOG) and specialized functional annotations such as biosynthetic gene cluster predictions using antiSMASH^40^.

Taxonomic classification of the genomic diversity in OMDB revealed a much improved representation of ocean microbial diversity compared to previous work across all taxonomic ranks (*e.g.*, n _species_ = 22,257; 7 times more than the second largest resource; Figure S2). To contextualize this expansion in diversity, we assessed species-level marker gene-based operational taxonomic units (mOTUs)^39^ and Average Nucleotide Identity (ANI) across OMDB, OceanDNA^30^ and GOMC^15^, allowing us to explore the prokaryotic diversity in each resource independently of taxonomic annotation. In total, genomes in OMDB mapped to 31,058 species-level clusters with 70.1% (n = 21,779) reported exclusively in OMDB (Figure S2). Beyond grouping genomes into species-level clusters, and as a distinguishing feature compared with other resources, we quantified relative species abundances across all samples enabling us to profile 45,497 species across the global ocean and perform quantitative analyses of microbial community composition along environmental gradients. Note that because the mOTUs profiler detects species directly from metagenomic reads, rather than requiring a reconstructed genome, it also captures species that are too rare or too poorly covered to be assembled into MAGs.

To facilitate interactive access to OMDB, we developed a web interface https://omdb.microbiomics.io as a centralized, globally contextualized resource for exploring ocean microbial genomes. As a starting point, the resource allows users to search, filter and download the genome, sample and study collections by tables and world maps. Furthermore, we made the 274,282 genomes, and over 508 million genes and proteins, searchable with the latest sequence search advances, including Metagraph^41^ and MMseqs2 expandable searches^42^, allowing users to rapidly retrieve sequence hits and visualize their geographic distribution (Figure S4). Apart from the rich interconnectivity between elements in the resource, all information is programmatically accessible via https://motus-api.microbiomics.io.

### Cross-study validation of temperature as primary driver of surface global ocean microbiome structure

To establish whether the patterns captured by the integrated OMDB data reflect generalizable biological trends rather than study-specific technical variation, we sought to reproduce and test the validity of a previously reported signal: the temperature-driven structuring of surface ocean microbial communities^1^.

We fitted Random Forest (RF) regressors to predict metadata variables from *Tara* abundance profiles (n = 371; epipelagic, prokaryotic size fraction) using 10-fold cross-validation. Models showed that community composition most strongly predicted temperature (R^2^ = 0.95), followed by nitrate/nitrite, silicate, and depth (Figure S5). We then extended this analysis using satellite-derived environmental variables, together with our curated in-situ temperature measurements to test whether these relationships remain at larger scales and across independent data sources. After applying a filtering criteria (epipelagic, prokaryotic-fraction, at least 10 samples per study with temperature data) and rarefying the abundance data (see Methods), the final dataset comprised 3,156 samples from 18 studies, with 11,827 detected mOTUs. Community composition continued to strongly predict temperature across this larger sample set: curated in-situ temperature (R^2^ = 0.95, n = 3,156) and satellite-derived temperature (R^2^ = 0.94, n = 1,905) were the most accurately predicted variables, followed by salinity (R^2^ = 0.84) and oxygen (R^2^ = 0.82). In contrast, ENA-reported temperature showed essentially no predictive power, highlighting the value of quality-controlled or remotely sensed data compared with raw metadata annotations (Figure S5; see Supplementary Material).

We tested whether the curated in-situ temperature prediction reflected a genuine microbiome-temperature relationship rather than study effects or spatial autocorrelation among proximate samples with similar temperatures. To this end, we applied several cross-validation strategies, including a strict geographic Leave-Neighbor-Out stratification between folds to ensure model generalization across larger distances (Figure S6). As output, we collected the R^2^ coefficients, Mean Absolute Error (MAE), and the importance of each mOTUs in the models (Figure 2A). Community composition remained a robust predictor of temperature (Figure S7) and showed well-supported correlation between actual and predicted temperature values (Figure 2B). Mean Pearson R^2^ was consistently high (∼0.6-0.9) with MAE remaining below 3 °C across the different split strategies (Figure 2C). The predictive power was maintained in temperate and tropical regions while a smaller number of held-out groups, concentrated at polar regions, showed reduced performance, consistent with these regions being comparatively undersampled, unevenly sampled across seasons, and environmentally distinct from the majority of the dataset (Figure S8).

**Figure 2.**
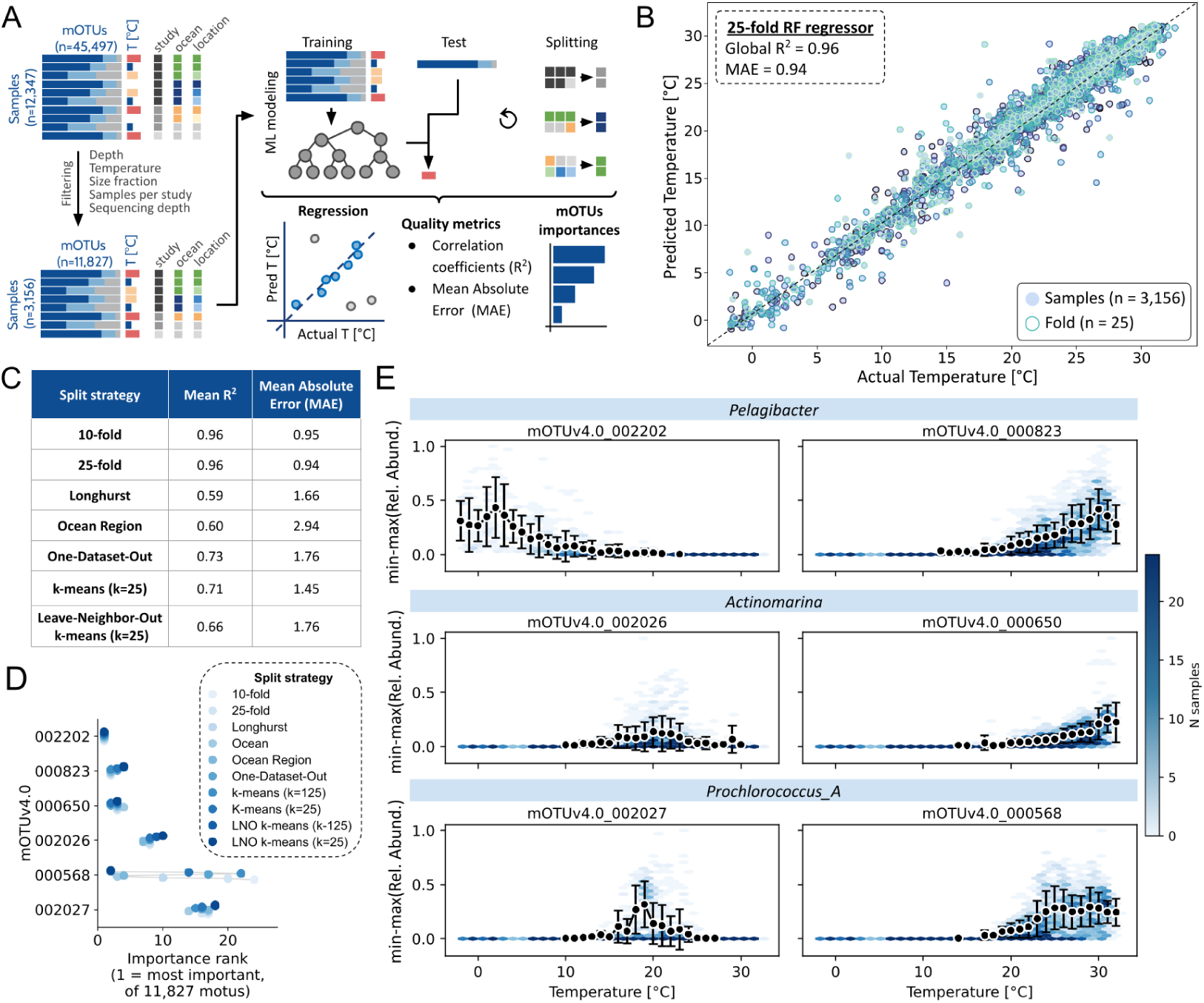
Machine learning reveals ocean microbiome composition as a robust predictor of temperature and identifies thermally partitioned sentinel taxa. **(A)** Schematic of the machine-learning workflow: mOTU relative abundance profiles were filtered, used to train regression models under multiple splitting strategies and evaluated by R^2^ and predicted-vs-actual agreement, and interrogated for mOTU feature importance. **(B)** Predicted versus observed temperature from 25-fold cross-validation of an RF regressor trained on the epipelagic dataset. Test-set sample predictions are shown and colored by fold. **(C)** Predictability of temperature under different stratified cross-validation strategies, designed to test generalization across studies, ocean regions, and larger geographic distances. **(D)** Feature importance rank of six representative mOTUs across all splitting strategies, illustrating consistently high importance regardless of splitting scheme. **(E)** Min-max scaled relative abundance of six highly explanatory mOTUs (genera *Pelagibacter*, *Actinomarina*, *Prochlorococcus_A*; top to bottom) versus temperature. Hexbins show sample density, including non-detections, on a shared colour scale per number of samples. Black points and error bars show mean ± SD per 1 °C bin, calculated only from samples where the mOTU was detected.

Having robustly reaffirmed the microbiome-temperature relationship, we sought to identify which taxa were driving the temperature predictability by analyzing RF feature importances. A *Pelagibacter* species (mOTUv4.0_002202) consistently emerged as the top individual predictor together with other model-informative species within *Pelagibacter*, *Actinomarina*, and *Prochlorococcus* genera (Figure 2D). These informative mOTUs exhibit distinct thermal optima in their abundance trajectories, suggesting thermal niche partitioning among congeneric species, illustrating how contrasting temperature associations among related taxa may contribute to broader thermal structuring of microbial communities (Figure 2E). Further analyses confirmed that predictive power did not rely solely on top-ranking taxa. An iterative feature drop-out analysis (Figure S9 and Figure S10), removing the mOTUs accounting for 90% of feature importance at each iteration, showed that performance remained stable across the first rounds, indicating redundancy among community members. In parallel, RF models were benchmarked against Ridge regression, which achieved comparable accuracy, but consistently underperformed in generalization and detection of informative mOTUs (Figure S11). Finally, models trained using only the four most explanatory mOTUs showed markedly degraded performance relative to the full model (Figure S12), confirming that robust prediction requires a broader ensemble of taxa rather than a handful of dominant predictors (see Supplementary Material for details).

Together, these results demonstrate that the temperature-community composition relationship generalizes robustly across independent datasets and spatial holdouts for the large majority of the ocean’s sampled regions, with additional data from polar environments expected to further improve predictive performance in these currently underrepresented areas.

### Realized thermal niche breadth predicts differential vulnerability of ocean microbial diversity to warming

We subsequently examined which community members may be the most vulnerable to additional warming and which are likely to endure. Because a taxon can only be detected within its current thermal tolerance range, we reasoned that species with narrower thermal niches would be more vulnerable as temperatures shift, whereas a taxon’s abundance would dictate the overall impact of its loss on the community. To test these relationships globally rather than focusing on a small number of well-studied taxa, we used the filtered dataset from the previous section.

For every mOTU, we fitted a smoothed abundance-temperature trajectory and used it to define their thermal niche breadth (Figure 3A) as the temperature range spanning the central 90% of the area under this trajectories, and yielding estimates for 7,021 mOTUs (Figure S13; see Methods). We then used these estimates to simulate the effect of warming under three IPCC Shared Socioeconomic Pathway (SSP) scenarios spanning a range of projected sea-surface temperature increases: SSP1-2.6 (+1 °C), SSP2-4.5 (+2 °C), and SSP5-8.5 (+3.5 °C)^6^. Richness declined steadily with warming: 89% of mOTUs remained detectable under SSP1-2.6, falling to 86% under SSP2-4.5 and 77% under SSP5-8.5 (Figure 3B). By contrast, the cumulative relative abundance retained by persisting mOTUs remained remarkably robust to the same scenarios, staying above 97% even under SSP5-8.5 (Figure 3B). Thus, even under the most severe warming scenario, ≥75% of detectable mOTUs are projected to persist, while richness declined roughly eightfold faster than abundance-weighted community structure (23% vs. 3% loss under SSP5-8.5; Figure 3C). The mOTUs projected to lose suitable habitat under warming therefore represent a disproportionately small share of total community relative abundance. This projected loss of richness should be interpreted cautiously, as richness is sensitive to detection limits and may overestimate the decline by failing to capture rare taxa.

**Figure 3.**
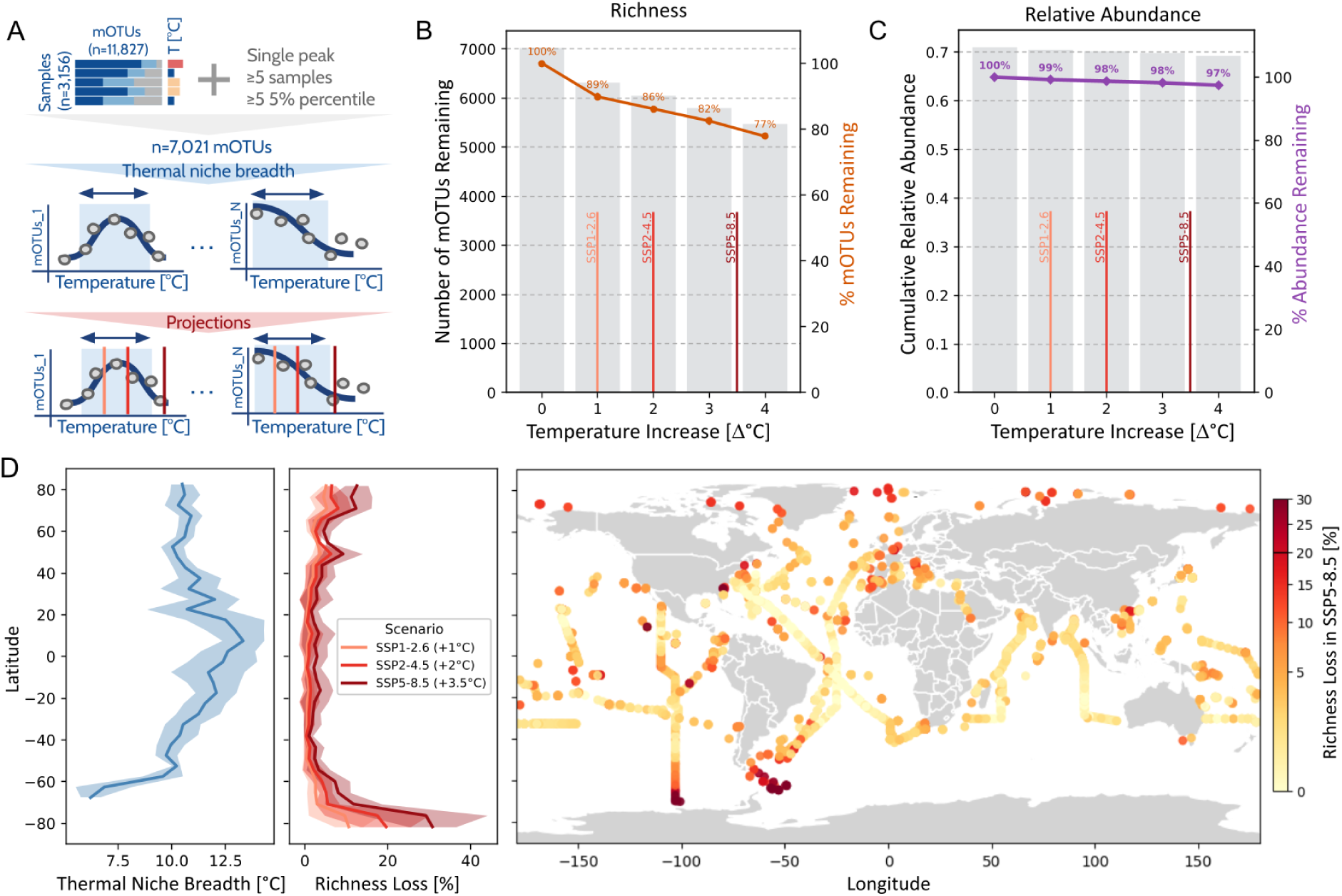
Thermal niche breadth reveals microbial communities susceptible to ocean warming. **(A)** Workflow: per-mOTU abundance-temperature trajectories are fitted and employed to define thermal niche breadth (blue box). Three IPCC scenarios are used to project them in relation to each species’ breadth. **(B)** Number and percentage of mOTUs retained as a function of simulated temperature increase (Δ°C), annotated at the IPCC scenarios (red-gradient dashed lines). **(C)** Cumulative relative abundance retained by the same persisting mOTUs, expressed as percentage of abundance remaining, under the same scenarios. Note the analyzed 7,021 mOTUs collectively account for ∼70% of total community relative abundance. **(D)** Left: mean thermal niche breadth by latitude. Middle: projected richness loss by latitude under the three SSP scenarios. Right: global map of projected fractional richness loss per sampling site under SSP5-8.5 (+3.5 °C), highlighting the Southern Ocean/Antarctic Peninsula as the strongest hotspot.

Both niche breadth and warming risk vary systematically with latitude. Mean thermal niche breadth was narrowest at high latitudes in both hemispheres and broadest across the tropical-to-temperate band (Figure 3D). Projected richness loss mirrored this pattern and was strongly scenario-dependent: under the mildest SSP1-2.6 scenario, richness loss remained below ∼10% across nearly all latitudes, whereas under the most severe SSP5-8.5 scenario losses increased toward the poles, exceeding 30% at the highest southern latitudes. Mapping these projections geographically under SSP5-8.5 (Figure 3D) identified the Southern Ocean and Antarctic Peninsula as the clearest hotspots of projected richness loss, with moderate losses (∼5-15%) across the Atlantic, Pacific, and Indian Oceans and little loss (<5%) in tropical and subtropical waters. Thermal niche breadth also varied with distance from the coast: mOTUs sampled within ∼1,500 km of the coastline had the broadest niches (∼12-14 °C), which narrowed to ∼9.4 °C in mid-ocean regions ∼3,500-3,700 km offshore (Figure S14). This pattern is consistent with greater thermal variability in coastal environments, driven by upwelling, mixing, and freshwater input, selecting for broader thermal tolerance than the more stable open ocean^43^.

Finally, mOTUs projected to be vulnerable under warming were significantly less likely to carry a formal species- and genus-level name in GTDB (Figure S15). In other words, mOTUs under the greatest thermal selective pressure are more often poorly characterized, and this bias may intensify with ocean warming, as poorly characterized taxa also tend to have the narrowest thermal niches.

### Discrete thermal-response clusters are underpinned by different genomic properties and physicochemical and thermostability adaptation of marker genes

To resolve whether the susceptibility gradient reflects continuous variation or discrete ecological strategies, we partitioned the abundance-temperature trajectories using unsupervised k-means clustering (Figure 4A). Among mOTUs with sufficient signal (n = 5,891; Figure S13; see Methods), a four-cluster solution (k = 4) provided the best separation of thermal-response patterns (Figure S16 and Figure S17), yielding four clusters ordered by ascending thermal optimum, from Cluster 1 (coldest) to Cluster 4 (warmest). Within the clusters 1 and 4, some mOTUs increased monotonically toward the coldest (n = 70) or warmest (n = 65) temperatures rather than showing an interior local maximum and they were analyzed separately from their parent clusters 1 and 4 as their thermal optimum likely lies outside the sampled range (Figure 4B; see Methods). Cold-associated mOTUs reached the highest relative abundances when detected (Figure 4C), indicating that these taxa are not simply rare across the dataset but can dominate local community composition within their realized thermal niche. The number of different clusters per sample peaked at mid-latitudes (∼30-50° in both hemispheres) and declined toward the poles and equator (Figure S18), suggesting that temperate zones harbor multiple thermal preferences, likely due to greater temperature variability or water-mass mixing, whereas polar and equatorial regions seem more homogeneous and dominated by a single thermal cluster. This should be interpreted cautiously, however, as OMDB sampling is skewed toward spring and summer months; we therefore lack the seasonal resolution to determine whether additional thermal niches exist in polar waters during ice-covered periods that are less represented in the dataset.

**Figure 4.**
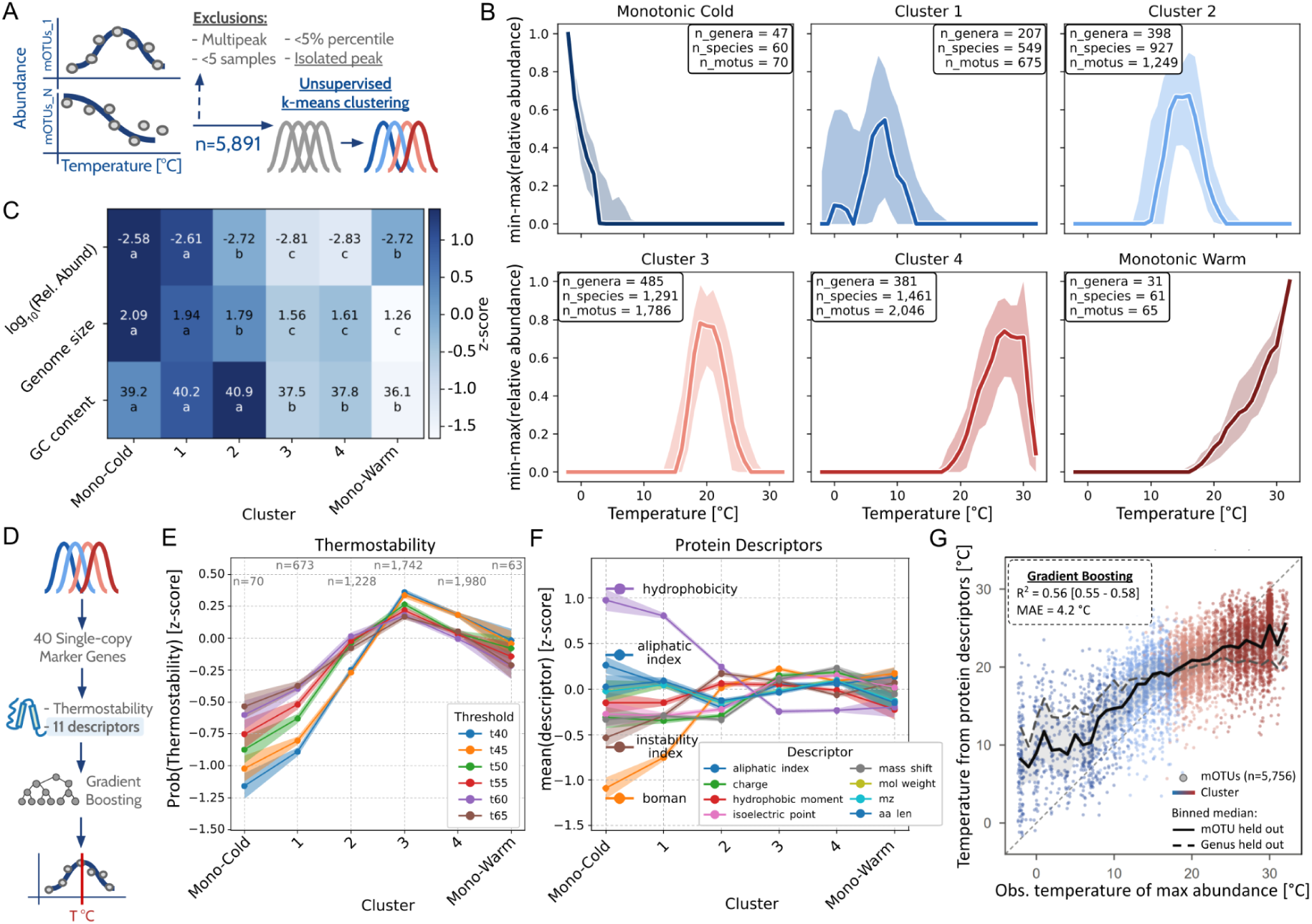
Genomic, proteomic, and physicochemical characterization of ecologically-defined thermal clusters. **(A)** Overview of clustering pipeline: per-mOTU abundance-versus-temperature trajectories were constructed, quality-filtered, and grouped by unsupervised k-means clustering. **(B)** Median (bold line) and interquartile range (shaded band) of normalized abundance trajectories for each thermal group. mOTUs with a monotonic (edge-truncated) trajectory were retained as separate groups rather than assigned to their classified cluster (1 and 4, respectively). **(C)** Mean relative abundance (when detected), Genome size (Mb), GC content (%) by thermal cluster; cell color indicates the row-wise z-score, and labels show the mean value and Tukey HSD post-hoc group (shared letters indicate no significant difference, α = 0.05). **(D)** Overview of protein-level analyses: we calculate thermostability and eleven descriptors from 40 SCMGs per mOTU and across clusters. With the protein descriptors we model temperature optima predictors. **(E)** Mean predicted thermostability (TemStaPro, z-scored) of single-copy marker genes (SCMGs) at six temperature thresholds (t40–t65) across thermal groups; shaded bands show ±SEM, and n indicates the number of mOTUs contributing predictions per group. **(F)** Mean physicochemical descriptor values (z-scored) of SCMG-encoded proteins across thermal groups; shaded bands show ±SEM. The number of mOTUs per cluster are shared with (E). **(G)** Observed temperature of maximum abundance versus the temperature predicted from the protein physicochemical descriptors. Each point corresponds to the prediction per mOTU, coloured by thermal cluster. The solid black line and grey band show the median and interquartile range of predictions per 1 °C bin. The dashed line shows the binned median when entire genera are held out of training (R^2^ = 0.26). R^2^, out-of-fold coefficient of determination, with a 95% bootstrap confidence interval (1,000 resamples of mOTUs); MAE, mean absolute error.

If these four ecological clusters reflect adaptive molecular differences, their genomes should carry distinguishing signatures (Figure 4D). Genome size and GC content declined from cold-to warm-associated groups (Figure 4C), consistent with genome streamlining and metabolic specialization under resource limitation^18,44,45^. To test whether this genome-level signal extends to finer molecular differences, we examined protein-level variation using 40 universal single-copy marker genes (SCMGs) with high species resolution, ensuring comparability across diverse taxa^46^. We then assessed SMCG thermostability using TemStaPro^47^, a protein embedding-based predictor, and found that it increased from the monotonic-cold cluster through cluster 3 before plateauing in the warmer clusters (Figure 4E). Among the 40 SCMGs, COG0012 (YchF), an ATPase involved in stress signaling and translational regulation, showed the strongest cold-to-warm increase (Spearman ρ = 0.94, p-value = 0.004; Figure S19), consistent with its reported role in thermal and oxidative stress responses^48^. The same gradient across 100 randomly sampled proteins per representative genome indicates broader thermal adaptation rather than a property of conserved SCMGs alone (Figure S20). To characterize thermal adaptation at finer amino acid sequence resolution, we next examined eleven physicochemical descriptors of SCMG-encoded proteins, all of which differed significantly across thermal clusters (Figure S21). Warm-associated clusters showed higher net charge, Boman index, and isoelectric point, consistent with a greater contribution of charged and polar residues, which is thought to stabilize proteins at higher temperatures and which is reduced in cold-adapted proteins^49^. In contrast, hydrophobicity and aliphatic index decreased toward warmer clusters, departing from signatures typically reported for thermophilic proteins^50^. Protein length (and hence molecular weight and m/z) decreased modestly from cold-to warm-associated clusters, mirroring genome streamlining (Figure 4F).

Given that these descriptors differ systematically across thermal clusters, we asked whether they predict where a mOTU reaches its maximum abundance in nature. Gradient-boosted models trained on the eleven descriptors of all 40 SCMGs predicted the temperature of maximum abundance of held-out mOTUs with R² = 0.56 (Figure 4G). Performance decreased but persisted when entire genera or families were withheld (R² = 0.26 and 0.20; Figure S22). The same descriptors assigned mOTUs to their thermal cluster with a balanced accuracy of 0.63, and 92% of mOTUs were placed in the correct or a neighbouring cluster (Figure S23). Hydrophobicity, Boman index and net charge contributed most to these predictions (Figure S24). Thus, although part of this signal reflects shared ancestry, the temperature at which a mOTU reaches its maximum abundance is partly imprinted in the combined physicochemistry of its core proteins

## Discussion

Efforts to link the genomic potential of the marine microbiome with global biogeochemical and ecological patterns have been hindered by the fragmentation of data and information across repositories and the primary literature, heterogeneous methods to reconstruct genomes from metagenomic samples, and inconsistent integration of metagenomic data with geographic and environmental metadata^51^. By integrating data from 209 studies through standardized genome reconstruction^14^, taxonomic and gene functional annotation, community taxonomic profiling and curation of geographic and environmental metadata, OMDB enables the research community to compare genomic potential, environmental distributions and ecological patterns across studies and at global scale.

Our results showing that photic-zone seawater temperature can be robustly predicted from microbial community composition independent from study, methodological and geographic effects generalize previous evidence^1^. Notably, our approach enables the data-driven identification of thermal niche-partioned congeneric members within *Pelagibacter*, *Actinomarina*, and *Prochlorococcus*. This was facilitated by the integrated abundance profiles provided by OMDB and represents a cultivation-independent route to studying ecological differentiation among closely related taxa. We note that temperature covaries with factors including light and nutrient availability, water mass structure and dissolved oxygen, for example. The observed associations therefore cannot distinguish direct effects of temperature from those of correlated environmental conditions or biological interactions. Disentangling these effects and extending similar analyses beyond the photic ocean layer remain important directions for future work.

While the scientific community has increasingly recognized the need to safeguard microbial biodiversity^9^, quantifying microbial vulnerability to climate change and the associated risk of extinction at a global scale remains conceptually and analytically difficult^52^. By estimating the realized thermal niches of species-level ocean microbial taxa, we asked in our work which members of the ocean microbiome are projected to encounter temperatures that exceed their present-day thermal niche boundaries. While crossing those boundaries does not necessarily imply exceeding physiological tolerance, it may indicate increased ecological vulnerability. Previous work examining the realized thermal niches of marine microorganisms has included comparisons of fundamental and realized niches among *Prochlorococcus* ecotypes and assessments of microbial responses to marine heatwaves^53,54^. However, to our knowledge, our study provides the first global, genome-resolved characterization of realized thermal niches across thousands of epipelagic marine microbial species-level taxa.

The asymmetry between richness and abundance loss is consistent with the uneven structure of microbial communities, in which a relatively small number of numerically dominant taxa account for most of the total observed relative abundance; the many individually rare taxa projected to lose suitable habitat therefore contribute comparatively little to total community abundance, regardless of their thermal niche breadth. Whether these numerically rare, more vulnerable taxa also differ systematically in effective population size, genetic diversity, or adaptive potential remains untested and will require dedicated population-genomic analyses. Geographically, the projected richness losses were concentrated in the Southern Ocean, around the Antarctic Peninsula and in the Arctic Ocean. Furthermore, the taxa projected to exceed the upper boundary of their realized thermal niches generally lack formal species names, highlighting that these losses may disproportionately affect poorly characterized taxa. However, these projections assume that present-day realized thermal niches remain fixed and do not account for physiological acclimation, evolutionary adaptation, or redistribution through dispersal, advection, and horizontal or vertical habitat shifts. Such redistribution may be especially important in the Arctic Ocean, where Atlantification can transport Atlantic populations northward while favoring temperate over Arctic specialists^55^. Moreover, the niche widths of rare taxa may be underestimated because of their lower detection frequencies, projected undetectability should not be interpreted as extinction, and changes in relative abundance do not directly represent changes in absolute abundance or biomass. Within these limitations, our findings add a species-resolved perspective to previous work suggesting that warming may cause particularly pronounced compositional changes in polar microbial communities^56^ and indicate that projected vulnerability is disproportionately concentrated among poorly characterized microbial lineages.

To resolve variation in present-day realized thermal niches, we clustered taxon-specific abundance-temperature trajectories into four discrete thermal-response groups and tested whether these patterns corresponded to molecular characteristics. Genome size and GC content generally declined from cold-to warm-associated clusters, consistent with genome streamlining under resource-limited and warm oligotrophic conditions^18,44,45^. At the protein level, thermostability predictions and physicochemical analyses revealed that warmer clusters shifted coherently toward features associated with known mechanisms of protein stabilization at higher temperatures^49,50^. When combined, these descriptors allowed us to predict directly from protein sequences the temperature at which each mOTU reaches its maximum *in situ* relative abundance. This concordance between sequence features and realized thermal niches suggests evolutionary adaptation and provides a basis for predicting the thermal preferences of uncultivated microorganisms from their genomes. Disentangling whether these patterns represent convergent thermal adaptation rather than shared evolutionary history will require further phylogenetic analyses. Also, thermostability was computationally predicted rather than measured directly, and the descriptor-based analyses were restricted to universal SCMGs. Extending the analyses across complete proteomes and validating the predicted structural and biochemical differences are therefore important directions for future work.

Ultimately, the integration of harmonized genomic, ecological, and environmental data within OMDB bridges molecular biology and global oceanography within a single resource. By linking geographic and ecological patterns (where organisms live, how abundant they are, how exposed they are to exceed their thermal niche boundaries) directly to the genomic and molecular properties, OMDB moves beyond a static genome repository to enable exploration of ocean microbial life across biological scales, from molecules to ecosystem. Together with the predictive frameworks presented here, spanning community-level machine learning to single-gene physicochemistry, we provide a baseline for monitoring, modeling, and predicting how the ocean microbiome, and the biogeochemical processes it drives, will respond to continued ocean warming.

## Limitations

The projected responses in our study do not account for physiological acclimation, ecological migration, or evolutionary adaptation that may occur before the projected mean ocean temperature increases are reached. Many marine microorganisms have short generation times and large population sizes, providing opportunities for evolutionary responses on timescales comparable to environmental change, including species-specific shifts in thermal tolerance^57^. Moreover, temporal fluctuations in ocean temperature may further accelerate microbial adaptation^58^. Modelling work further suggests that short generation times can lead to genetic adaptation on timescales comparable to environmental change^59^. Our estimates are also less certain for sparsely detected taxa with poorly constrained realized thermal niche boundaries. Furthermore, we acknowledge that the cumulative relative abundances of our dataset are influenced by uneven geographic sampling rather than an area-, volume-, and biomass-weighted representation of the global ocean microbiome. Resolving physiological and adaptive responses would benefit from metatranscriptomic, metaproteomic, and genomic long-term time series, and representative population size estimates from using quantitative methods that estimate taxon-specific cell concentrations and biomass^60,61^. Nonetheless, our work provides a data-driven, taxon-resolved framework for identifying marine microorganisms under potential ecological vulnerability, and a quantitative baseline at global scale. Linking these projections to Earth system models, and testing whether expanding generalists retain the functional capacity of the specialists they might replace remain important challenges.

## Methods

### Genomic data collection

A total of 209 publicly available studies, published up to December 2023, were gathered from selected marine metagenomics literature and matched with their respective BioProject identifiers from the ENA. To ensure environmental relevance, and consider only samples from projects containing different environmental samples, this initial dataset was filtered further based on ENA metadata attributes and by geographic location constraints using the Python library global-land-mask v1.0.0. Reference genomes were simultaneously downloaded from GTDB and ProGenomes 3 genome collections^62,63^, excluding non-marine labelled entries and metagenome-assembled genomes (MAGs), using the NCBI Datasets command-line tool^64^. In addition, 70 genomes were retrieved from the JGI Genome Portal. After filtering for assembly quality (’complete genome’ or ‘representative genome’ status), 13,730 non-MAGs genomes were integrated into the local database, as done for mOTUs-db^39^. As a quality control measure to ensure dataset comprehensiveness, we cross-referenced our collection with samples included in other established reference databases. These included MarDB and MarRef from the Marine Metagenomics Portal (https://sfb.mmp2.sigma2.no/) and Planet Microbe (https://www.planetmicrobe.org/) (both accessed in April 2024). Through this comparative analysis, we confirmed that all samples from these external repositories were in fact encapsulated within our finalized Ocean Microbiomics Database (OMDB) framework.

### Sequencing data processing

For each study collected, raw read data were downloaded from ENA, and metagenomic data processing was performed as described in Paoli et al.^14^. Briefly, sequencing raw reads were filtered using BBMap (v.38.06) by removing sequencing adapters from the reads, filtering out reads that mapped to quality control sequences (PhiX library), and discarding low quality reads using the parameters t*rimq = 14, maq = 20, maxns = 1*, and *minlength = 45*. The resulting quality-controlled reads were then merged using BBMerge (v.38.06) (*minoverlap = 16*), resulting in merged, unmerged paired-end and single-end reads. Alternatively, read sets from Tara expeditions and from samples that required >2TB of RAM in the subsequent assembling step were normalized with bbnorm.sh *target = 40* and *mindepth = 0*.

### MAG reconstruction, quality assessment and taxonomic profiling

The SPAdes assembler^23^ (v3.11-v3.15) in metagenomic mode was run to assemble these reads into scaffolded contigs (hereafter scaffolds) To generate multi-sample coverage information for binning scaffolds, we grouped samples into groups of size ≥50 based on the environment and study from which they were derived. Quality-controlled metagenomic reads from each sample were mapped to scaffolds of at least 1000 bp from all samples within the same group. The mapping was performed using BWA (v.0.7.17-r1188)^65^ and allowing the reads to map at secondary sites (with the *-a* flag). Alignments were then filtered to be ≥45 bases in length, with an identity of ≥97% and covering ≥80% of the read sequence. The BAM output files were processed using the *jgi_summarize_bam_contig_depths* script of MetaBAT2 (v.2.15)^66^ to provide within- and between-sample coverages for each scaffold. The scaffolds were binned by running MetaBAT 2 on all samples individually with parameters *--minContig 2000* and *--maxEdges 500* for increased sensitivity. The quality of each metagenomic bin was evaluated using both the ‘lineage workflow’ of CheckM^67^ (v.1.1.3) and Anvi’o^68^ (v.7.1). Quality labels were assigned to each genome according to completeness and contamination values following community standards^69^. Of the 260,552 MAGs in OMDB, 57,967 (22.2%) were of high quality (≥ 90% completeness and ≤ 5% contamination), 91,706 (35.2%) of good quality (≥ 70% and ≤ 10%), 79,741 (30.6%) of medium quality (≥ 50% and ≤ 10%) and 31,138 (12.0%) of fair quality (≤ 90% completeness or ≥ 10% contamination). An additional 13,730 reference genomes from isolates were included without quality labelling (Figure S2). MAGs with CheckM completeness ≥50% and contamination ≤10% or Anvi’o completion ≥50% and redundancy ≤10% were taxonomically annotated using GTDB-Tk^70^ (v.2.4.0) with the default parameters against the GTDB R226 release^71^.

In parallel to genome reconstruction, quality-controlled metagenomic reads from all samples were taxonomically profiled using the mOTUs database^39^ (v4.0; default parameters).

### Gene sequences catalog generation

Gene sequences were predicted using Prodigal^72^ (v2.6.3) with the parameters *-c -q -m -p single*. Gene and protein sequences (n = 508,832,278) predicted across all 274,282 genomes were aggregated into non-redundant nucleotide (NT) and amino acid (AA) catalogs. Redundancy was reduced using the MMseqs2^42^ clustering module at multiple thresholds. The NT catalog was clustered at 100% (NR100; n = 325,384,975) and 95% (NR95; n = 103,044,829) identity. The AA catalog was clustered at 100% (NR100; n = 249,518,434), 50% (NR50; n = 28,862,112), and 30% (NR30; n = 18,342,415) identity, providing representative sequence sets at varying resolution for downstream mining and search.

In addition, single-copy marker genes were extracted using fetchMGs (v1.2), with parameters -m extraction *-v -i*, yielding sequences for 40 genes identified by their COG number for each genome^46^.

### Genome annotation and genome-resolved information

Each genome was annotated using standardized procedures to ensure consistent and reproducible functional annotations. Ribosomal RNA genes were predicted using barrnap (v0.9, https://github.com/tseemann/barrnap) run separately with bacterial and archaeal models (*--kingdom bac / --kingdom arc*). Transfer RNA and transfer-messenger RNA genes were identified using ARAGORN^73^ (v1.2.41) with the bacterial/archaeal genetic code (*-l -gc11*). Protein-coding genes were functionally annotated against Pfam^74^ (v37.1) using pyHMMER^75^ (v0.11.2) with gathering-threshold bit-score cutoffs, and against the eggNOG (v5.0.2) orthology database using eggNOG-mapper^76^ (v2.1.7) with DIAMOND^77^ alignment in sensitive mode (*-m diamond --sensmode more-sensitive*). KEGG orthologs and pathways were annotated separately by DIAMOND^77^ (v2.0.15.153) alignment against the KEGG database (April 2022) with *--id 25 --query-cover 70 --subject-cover 70 --top 20 --more-sensitive* parameters. Identification and annotation of biosynthesis gene clusters was performed using antiSMASH^40^ (v6.1.1).

### Web application

The OMDB web interface (https://omdb.microbiomics.io) was implemented using the Flask (v2.3.2) Python web framework, with sample, genome, and taxonomic metadata served from an SQLite using Flask-SQLAlchemy (v3.0.3) database. Interactive geographic visualizations were built using Leaflet.js (v1.7.1). The interface supports browsing, filtering, and downloading genome, sample, and study collections via tabular and map-based views (Figure S4). The data contained in OMDB can be accessed programmatically through https://motus-api.microbiomics.io.

Rapid sequence-based querying of the OMDB catalogs is supported through multiple search backends. Exact/near-exact matches can be retrieved via standard BLASTN^78^ (BLAST+ v2.15.0) against the genome and the gene catalogs. We also integrated Metagraph^41^ for indexed, petabase-scale retrieval against the full non-redundant nucleotide gene catalog (OMDv2.0_NT_GENO_NR), queried via the Metagraph HTTP API. Expandable protein searches use MMseqs2^42^ via its expanded-cluster search workflow (mmseqs search → expandaln → align → convertalis), allowing retrieval of homologous hits, with results visualized by geographic and taxonomic distribution (Figure S4).

### Genome-resolved resources comparative

For quality comparisons with GOMC and OceanDNA, we used CheckM metrics. GOMC quality metrics were available for only 24,195 representative genomes, so we ran the same CheckM pipeline used for OMDB on the GOMC-specific MAGs included in our comparison. For the per taxa-level comparisons, all genomes from OMDv1, OceanDNA and GOMC were re-annotated using GTDB-Tk (v.2.4.0)^70^ with the default parameters against the GTDB R226 release^71^. For mOTUs-level comparatives, GOMC and OceanDNA were mapped onto existing mOTU species clusters. Finally, to compare genome-level redundancy and uniqueness across OMDB, GOMC, and OceanDNA independently of each resource’s own taxonomic assignments, pairwise average nucleotide identity (ANI) was computed for the combined genome set from the three databases using skani triangle with default parameters. Genomes were grouped into dereplicated clusters via single-linkage transitive clustering on the resulting pairwise ANI matrix: any two genomes with ANI ≥ 99% were considered to belong to the same cluster.

### Predicting temperature from microbial abundance profiles

Random Forest (n_estimators = 100) was imported from scikit-learn (v1.6.1) and trained to predict in situ temperature from mOTU relative abundance profiles.

- <u>Sample selection and rarefaction</u>: We first selected 3,630 samples fitting the following criteria: i) epipelagic layer (surface down to 200 meters); ii) prokaryotic size fraction; iii) salinity > 30 psu (excluding *e.g.*, samples from the Baltic sea); and iv) minimum 10 samples per study. Selected samples were rarefied to ∼ 1,000 reads using skbio.stats.subsample_counts (v0.6.3), chosen to balance sample retention against total and per-sample mOTU richness and dataset coverage. mOTUs with zero total counts across samples after rarefaction were dropped; no additional prevalence or minimum-abundance filter was applied beyond this, yielding a final dataset of 3,156 samples and 11,827 mOTUs.
- <u>Splitting strategies</u>: Model generalization was evaluated using standard 10-fold and 25-fold cross-validation, leave-one-dataset-out, leave-one-Longhurst-province-out, leave-one-ocean/ocean-region-out, and spatially aware k-means splits (k = 25 and k = 125 geographic clusters, fit on sample latitude/longitude with scikit KMeans [v1.6.1]). The latter were run both as conventional k-fold cross-validation and as a stricter “leave-neighbor-out” (LNO) scheme, in which clusters within a minimum centroid distance of 5,500 km (great-circle distance via geopy.distance.geodesic [v2.4.1]) of the held-out test cluster were also excluded from training (Figure S6) to guard against inflated performance from spatial autocorrelation between nearby, thermally similar samples.
- <u>Robustness and benchmarking</u>: An iterative feature drop-out analysis (Figures S9 and Figure S10) removed, after each training round, the mOTUs collectively explaining 90% of feature importance, and repeated model training for 20 iterations to assess predictive redundancy. RF performance was additionally benchmarked against Ridge regression (features standardized with StandardScaler prior to fitting) using identical splits (Figure S11), including a reduced-feature comparison using only the four most explanatory mOTUs (Figure S12). The Ridge regularization parameter (alpha) was selected via a regularization-path search over 30 log-spaced values between 1e-5 and 1e6 under 10-fold cross-validation. Although the cross-validated optimum was alpha = 2212.2 (R^2^ = 0.968), alpha = 1 was used for all reported comparisons to preserve coefficient interpretability rather than to maximize cross-validated R^2^ (see Supplementary Material).
- <u>Evaluation</u>: Performance was assessed via Pearson and Spearman R^2^ and mean absolute error (MAE, °C) between predicted and observed temperature on held-out test samples. Importances within each model were calculated as the average reduction in the regression-tree splitting criterion attributable to each feature, averaged across all trees in the forest.

### mOTU thermal trajectory construction and quality filtering

To characterize the thermal preference of individual mOTUs, we constructed per-mOTU abundance trajectories across a temperature gradient and applied a series of quality filters to retain only mOTUs with a confidently resolved thermal optimum for downstream clustering. Starting with the same rarefied table produced for the RF regressor analysis (11,827 mOTUs along 3,156 epipelagic and prokaryotic size fraction samples with temperature available), samples were first binned into 1 °C temperature intervals based on their associated in-situ temperature measurement. mOTUs with fewer than 5 total non-zero detections across the full dataset were excluded prior to binning. For each remaining mOTU, mean abundance (including non-detections as zero) was computed per temperature bin. To ensure bin-level reliability, a bin’s value was retained only if supported by at least 3 samples with non-zero detection; bins failing this threshold were set to zero. Subsequently, binned trajectories were smoothed using a Savitzky-Golay filter (window = 5 bins, polynomial order = 2, boundary mode = nearest to avoid edge extrapolation artifacts), clipped at zero to remove residual smoothing overshoot, and min-max normalized per mOTU to a 0-1 scale. mOTUs were then excluded from the clustering table if they met any of the following criteria, evaluated on the smoothed, normalized trajectories unless otherwise noted:

- **Below threshold**: maximum raw binned abundance fell below an empirically derived cutoff (5th percentile of the maximum abundance distribution among mOTUs with non-zero signal), indicating insufficiently reliable signal.
- **Isolated peak**: the entire non-zero signal was confined to a short (≤2-bin), zero-flanked run (≥2 bins of zero on both sides), consistent with a spurious or non-generalizable detection rather than a genuine thermal response.
- **Multipeak**: the trajectory exhibited ≥2 robust local maxima (minimum height 0.5 on the normalized scale, minimum prominence 0.15, separated by ≥10 °C), indicating an ambiguous or compound thermal signal not attributable to a single optimum.
- **Monotonic to boundary**: the trajectory showed no interior local maximum and instead increased monotonically toward one edge of the sampled temperature range (peak value ≥0.5 at the first or last bin), indicating the true thermal optimum likely lies outside the sampled range. These mOTUs were further classified as monotonic_cold or monotonic_warm according to which boundary they approached.
- **Flat trajectory**: the normalized trajectory showed a dynamic range (max - min) below 0.1, indicating no meaningful abundance variation across the temperature gradient despite passing the below-threshold filter on raw magnitude. This category was introduced to capture mOTUs with sufficient raw abundance but no shape-based thermal signal. This pattern related to rare mOTUs whose detection frequency, but not detection magnitude, varied with temperature.

Thermal niche breadth was defined, for each mOTU passing the filters above, as the number of 1 °C temperature bins spanning the central 90% of the cumulative area under its smoothed, normalized abundance trajectory (*i.e.*, excluding the outermost 5% of trajectory area on each side). One mOTU (mOTUv4.0_000793) was manually reclassified from an automatically assigned category to multipeak following visual inspection, as its raw detection pattern showed two clearly separated thermal associations that narrowly fell below the automated peak-prominence threshold.

### Simulating a temperature shift and its ecological impact

For a simulated warming shift of Δ°C, discretized into 1 °C bins, a mOTU was defined to persist (remain detectable) if at least one bin of its thermal niche remained after shifting its lower edge upward by that many bins; equivalently, a mOTU was projected to be lost once the simulated shift exceeded its (unshifted) niche breadth. This criterion was applied dataset-wide at three IPCC Shared Socioeconomic Pathway (SSP) warming levels: SSP1-2.6 (+1 °C), SSP2-4.5 (+2 °C), and SSP5-8.5 (+3.5 °C, evaluated at the nearest 1 °C bin)^6^. Two quantities were reported at the whole-dataset level for each scenario: (i) the number and percentage of all analysis-eligible mOTUs projected to remain detectable, and (ii) the cumulative relative abundance of persisting mOTUs — defined as the sum, over the surviving mOTU set, of each mOTU’s mean relative abundance across all samples (including samples where it was absent) — expressed both in absolute terms and as a percentage.

Latitudinal trends (Figure 3D) were obtained by binning samples into 5° latitude bands and reporting the mean ± SD of the per-sample loss fraction (or, for thermal niche breadth, the abundance-weighted mean breadth of locally present mOTUs) within each band. Distance to coastline (Figure S14) for each sample was computed as the geodesic distance from the sample’s coordinates to the nearest point on the global coastline boundary, derived from the Natural Earth 1:110m physical land vector dataset (ne_110m_land), using geopandas (v1.0.1).

GTDB species- and genus-level names were classified as genuinely resolved or as unresolved placeholders; a name was treated as unresolved if it was missing, labeled “Unclassified,” prefixed “Unknown,” or matched GTDB’s placeholder-code naming convention (alphanumeric codes containing digits, or all-caps tokens longer than three characters, rather than a capitalized Latin binomial; *e.g.*, “UBA1014,” “GCA-002686595,” “SMXP01”). For each SSP scenario, the rate of species-level (and, separately, genus-level) resolution among mOTUs projected to be lost was compared against the rate among mOTUs projected to survive using a two-sided Fisher’s exact test on the resulting 2×2 contingency table (lost/survived × resolved/unresolved); p-values across the three scenarios were adjusted for multiple comparisons using the Benjamini-Hochberg procedure (Figure S15).

### Unsupervised clustering of abundance response in a temperature gradient

The retained mOTUs and their trajectories were clustered and evaluated using scikit-learn (v1.6.1) k-means and clustering performance metrics. The number of clusters (k) was selected by jointly evaluating six complementary metrics across k = 2-19 and using all possible metrics described to be informative in similar contexts^79^: inertia (elbow method), silhouette score, Davies-Bouldin index, Calinski-Harabasz index, bootstrap stability (mean adjusted Rand index across 20 bootstrap resamples of the input trajectories), and prediction strength. k = 4 was selected as the value at or near the elbow/optimum of the inertia, silhouette, Davies-Bouldin, and Calinski-Harabasz curves, within the range where bootstrap stability remained high (mean ARI > 0.9) and where prediction strength first exceeded the conventional reproducibility threshold of 0.8 (Figure S16). A secondary local maximum in prediction strength near k = 9 did not correspond to additional, visually distinct trajectory shapes and was not considered further (Figure S16 and Figure S17).

Cluster identifiers were reordered post hoc by ascending median peak temperature (Cluster 1 = coldest, Cluster 4 = warmest) for interpretability. Within the two boundary clusters (1 and 4), the “monotonic” mOTUs identified during shape classification (above) were displayed separately from the remaining, interior-peaked majority of their parent cluster, but were not treated as separate clusters for the k-means solution itself.

### Cluster-level genomic and abundance comparisons

Genome size, GC content, and mean relative abundance (log10-transformed, calculated per mOTU only across the samples in which it was detected) were each compared across the four thermal clusters (and, where shown separately, the two monotonic subsets) using one-way ANOVA followed by Tukey’s HSD post-hoc test. The same one-way-ANOVA-plus-Tukey-HSD approach was used to compare each of the eleven physicochemical descriptors (below) across clusters, and Kruskal-Wallis tests (with Benjamini-Hochberg-corrected p-values where multiple features/COGs were tested jointly) were used where normality of the underlying feature distributions was not assumed (Figure S21).

### Predicted protein thermostability

Predicted thermostability of SCMG-encoded proteins was assessed using TemStaPro (v0.2.6)^47^, a sequence-embedding-based classifier that assigns each protein a predicted-stability probability at six discrete melting-temperature thresholds (40, 45, 50, 55, 60, and 65° C). Predictions were generated for (i) the SCMG sequences described above, restricted to mOTUs represented by at least one MAG in OMDB (n = 5,756 mOTUs with an assembled genome in OMDB), and (ii) a background set of 100 randomly sampled protein sequences per mOTU representative genome, to test whether any thermostability gradient was specific to marker genes or detectable genome-wide (Figure S20). Per-mOTU scores at each threshold were aggregated (mean) and z-scored for cross-cluster comparison. Association between predicted thermostability and thermal-cluster assignment was tested per COG and per threshold using one-way ANOVA, and COGs were ranked by the strength of this association (η²) and, separately, by the monotonicity of their cold-to-warm trend (Spearman ρ between cluster order and mean thermostability), to identify individual marker genes such as COG0012 (YchF) with the clearest cluster-thermostability relationship (Figure S19).

### Prediction of thermal niches from core-protein physicochemistry

- <u>Features:</u> For each of the 40 SCMGs in every genome belonging to 5,756 mOTUs), we computed eleven physicochemical descriptors of the encoded protein with the Python package peptides (v0.3.4): aliphatic index, Boman index, net charge, hydrophobic moment, hydrophobicity, instability index, isoelectric point, mass shift, molecular weight, m/z and length. We also computed the GC content of each gene. Genomes of the same mOTU carried near-identical SCMG sequences (median within-mOTU SD ≤ 6% of the between-genome SD for every descriptor), so features were summarised per mOTU as the median across its genomes for each SCMG.
- <u>Targets</u>: The regression target was the temperature bin (1 °C) at which each mOTU reached its maximum relative abundance. The classification targets were the four trajectory clusters and a six-class version in which the cold- and warm-monotonic subsets were treated separately.
- <u>Models</u>: We used histogram-based gradient boosting (scikit-learn v1.6.1; HistGradientBoostingRegressor and HistGradientBoostingClassifier) with fixed hyperparameters: 300 iterations, learning rate 0.05, at most 15 leaf nodes, at least 10 samples per leaf, L2 regularisation 1.0. Classifiers used balanced class weights, missing values were handled natively. Four feature sets were compared: 1) genome size and genome GC content only (baseline); 2) each SCMG separately, i.e. 40 independent models trained on the 11 descriptors of one SCMG each; 3) an ensemble of these 40 single-SCMG models, obtained by averaging their out-of-fold predictions for each mOTU (for classification, averaged class probabilities); a single model trained on the descriptors of all 40 SCMGs jointly (440 descriptors), which can combine information across genes. Each set was fitted with protein descriptors alone and with gene and genome GC content added.
- <u>Cross-validation</u>: Performance was assessed on out-of-fold predictions from 5-fold cross-validation under three schemes of increasing phylogenetic independence: i) mOTU held out (random folds of mOTUs, stratified by class for classification); ii) Genus held out (all mOTUs of a GTDB genus assigned to the same fold); iii) Family held out (all mOTUs of a GTDB family assigned to the same fold). For grouped regression, groups were assigned to folds to balance fold sizes; for grouped classification, StratifiedGroupKFold was used. Each mOTU’s genus and family were taken from its representative genome (majority assignment across its genomes if none was designated). Unclassified ranks were set to the nearest classified parent, so related unclassified mOTUs were kept in the same fold. Within each target and scheme all models shared identical folds.
- <u>Performance metrics</u>: Regression performance was quantified as the out-of-fold coefficient of determination (R^2^ = 1 − SS_res/SS_tot) and the mean absolute error. Classification performance was quantified as balanced accuracy, the mean recall across classes, which is insensitive to class size (chance = 1/number of classes). Because the thermal clusters form an ordered cold-to-warm series, a mOTU assigned to a neighbouring cluster (e.g. cluster 2 instead of cluster 3) represents a smaller error than one assigned to the opposite end of the gradient (e.g. cluster 4 instead of cluster 1). Balanced accuracy does not distinguish these cases, so we also report the fraction of mOTUs assigned to either the correct or a directly neighbouring cluster. 95% confidence intervals were obtained by bootstrap resampling of mOTUs (1,000 resamples for R², 300 for balanced accuracy).
- <u>Significance and feature importance</u>: For the joint models, significance was assessed against a null distribution from 10 label permutations run on the same folds, and is reported as the z-score of the observed performance relative to that null. Feature importance was estimated by grouped permutation on each test fold: each descriptor was permuted simultaneously across all SCMGs and, separately, all descriptors of each SCMG. Importance was recorded as the resulting loss in R^2^ or balanced accuracy, averaged over the five folds.
- <u>Within-genus analysis</u>: To test whether predictions discriminate between closely related mOTUs, we took genera with ≥ 5 mOTUs (240 genera, 4,674 mOTUs). For each, we correlated (Pearson) the observed and predicted deviations from the genus mean. A null distribution was obtained by shuffling predictions within genera 200 times.
- <u>Ecological covariates</u>: To evaluate whether the protein signal could be explained by covariates other than protein composition, we fitted the same model on two ecological covariates (log_10_ mean relative abundance and log_10_ number of recovered genomes) and two genome-level covariates (genome size and genome GC content). For the temperature of maximum abundance, these models used the same folds as the protein models. The out-of-fold predictions of the two models were then combined by linear regression, itself fitted within the same folds. The independent contribution of protein descriptors was quantified as the partial correlation between observed values and protein predictions, controlling for the covariate predictions.

## Acknowledgments

We thank Jonas Schiller and Dominic Eriksson for supporting fraction size metadata curation and Guillem Salazar for their contributions to study collection. We thank Nathan Beech for discussions on the warming-projection analysis. We thank Milot Mirdita and Martin Steinegger for their support in the integration of MMSeqs2 for sequence searches. We would like to thank all members of the Sunagawa, Christian von Mering, and Peer Bork labs for usability testing and feedback on the web interface. We would also like to thank the people we met at conferences and our other collaborators who generously provided thoughtful feedback throughout the various stages of OMDB’s development. We are also grateful to the research community for providing the data on which this resource is built. Finally, we are grateful to ETH IT services and HPC facilities for granting access to the EULER high performance cluster and providing web hosting expertise and support.

S.M.V. acknowledges funding from the Human Frontier Science Program through the fellowship LT0050/2023-L (https://doi.org/10.52044/HFSP.LT00502023-L.pc.gr.171942). T.P. acknowledges funding from the NOMIS Foundation through the NOMIS-ETH Postdoctoral Fellowship program. This work was funded by the Swiss National Science Foundation project grant 205320_215395 and the Swiss National Centre of Competence in Research (NCCR) “Microbiomes” grants 180575 and 225148 to S.S. This is publication number X of *Tara* Oceans and publication number Y of *Tara* Pacific.

## Competing interests

Authors declare no competing interests.

## Supplementary Material

### Supplementary Information

#### Modeling temperature prediction from abundances using Random Forest on *Tara* and satellite derived metadata

The Random Forest model predicted environmental variables in test samples (10-fold cross-validation) with the highest performance for temperature (R^2^ = 0.95, mean Pearson R^2^ across the folds) followed by silicate, nitrate/nitrite, and depth (Figure S5). This was followed by nitrate/nitrite (R^2^ = 0.81), silicate (R^2^ = 0.80), and depth (R^2^ = 0.79). These associations were independently supported by PERMANOVA performed on the same community abundance data, which yielded significant results for every factor tested (all p = 0.001; pseudo-F ranging from 1.85 for depth to 3.97 for N*).

To determine whether this relationship remains consistent at a larger scale and across independent data sources, we extended the analysis using satellite-derived environmental variables from Copernicus Marine Service, together with our curated in-situ temperature measurements. Community composition continued to strongly predict temperature across this larger and more heterogeneous sample set (Figure S5): curated in-situ temperature (R^2^ = 0.95, n = 3,163) and satellite-derived temperature (R^2^ = 0.94, n = 1,905) were the most accurately predicted variables, followed by salinity (R^2^ = 0.84), oxygen (R^2^ = 0.82), phytoplankton biomass (R^2^ = 0.80), nitrate and phosphate (both R^2^ = 0.78), and pH (R^2^ = 0.72; all n = 1,905). In contrast, metadata-derived temperature annotations showed essentially no predictive power (mean R^2^ ≈ 0, n = 999), highlighting the value of quality-controlled or remotely sensed data compared with raw metadata annotations. A complementary Mantel test comparing community dissimilarity with the dissimilarity of each environmental variable further confirmed these associations (all p = 0.001). Satellite-derived temperature showed the strongest correlation (R^2^ = 0.57), followed by curated in-situ temperature (R^2^ = 0.54), oxygen (R^2^ = 0.44), nitrate (R^2^ = 0.33), phosphate (R^2^ = 0.33), phytoplankton (R^2^ = 0.32), pH (R^2^ = 0.28), and salinity (R^2^ = 0.19). Metadata-derived temperature again showed the weakest association (R^2^ = 0.17).

#### Further evaluation of the models predictability by mOTUs iterative drop-outs

Because the strongest individual predictors identified in the main analysis were a small number of taxa with highly constrained thermal preferences (Figure 2E), we asked whether the models predictive power genuinely depended on this narrow group of sentinel species, or whether these taxa were simply the most conspicuous representatives of a broader, redundant thermal signal distributed throughout the community. This distinction is important for assessing the generalizability of our findings: if predictive performance collapsed after removing these few species, the temperature-community association would be fragile and dependent on a handful of cosmopolitan taxa, weakening its interpretation as a general ecological pattern rather than an artifact of a few dominant, well-sampled species. We therefore performed an iterative feature drop-out analysis to directly assess the depth and redundancy of the temperature signal encoded across the full mOTU community.

At each iteration N+1, we removed all mOTUs that collectively accounted for up to 90% of the regressor’s variance at iteration N, retrained the model using the remaining features, and repeated this process for 20 iterations across each splitting strategy (Figure S9). Across splitting strategies, the number of mOTUs required to reach the 90% variance threshold increased during iterations 5-10, reaching up to ∼1,000 mOTUs (Figure S9A), before declining as the pool of informative features was progressively depleted. In contrast, the total number of available features decreased steadily and approximately linearly across iterations as increasingly redundant taxa were removed (Figure S9B). Model performance, measured by R^2^ and MAE, remained relatively stable during the first several drop-out rounds, demonstrating substantial predictive redundancy among mOTUs. Thus, many taxa appear to encode overlapping thermal information, allowing the loss of individual predictors, or even the highest-explaining subset, to be compensated for by other members of the community.

However, after approximately 10-12 iterations, model performance deteriorated sharply and consistently across all splitting strategies (Figure S9D and Figure S9D). R^2^ declined toward zero while MAE increased several-fold, indicating that the redundancy of the thermal signal is finite. Once the broader pool of thermally informative taxa was exhausted, the residual community could no longer maintain predictive performance. Notably, the space-aware splitting strategies that showed the weakest baseline performance (*e.g.*, ocean and ocean_region) also exhibited the steepest early increases in error. This suggests that generalization across ocean basins depends disproportionately on the broader redundant pool of thermally informative taxa rather than on a small, fixed set of sentinel taxa.

Tracking the variance explained by the single most important mOTU at each iteration provided complementary evidence for this interpretation (Figure S10). Following removal of the initial top-explaining set, the maximum single-feature importance decreased from approximately 0.4-0.55 to below 0.1 within the first 3-5 iterations across all splitting strategies. It then increased again toward the final iterations as the feature pool became progressively smaller, such that individual remaining mOTUs necessarily accounted for a larger fraction of the much-reduced explainable variance. This late increase therefore reflects feature scarcity rather than a recovery of strong individual predictive power.

Together, these results indicate that the sentinel taxa highlighted in the main results are best interpreted as clear and illustrative examples of a temperature signal that is distributed redundantly across a much larger set of community members, rather than as the sole drivers of model performance.

#### Employing Ridge Regressors as alternative modeling approach

To further validate the Random Forest results, we benchmarked model performance against Ridge regression using identical data splits. Using the 10-fold cross-validation strategy we first optimized the Ridge regularization parameter (alpha) via a regularization path search (Figure S11A). Mean cross-validated R^2^ peaked at alpha = 2212.2 (R^2^ = 0.968). However, this alpha value corresponds to strong shrinkage, pushing coefficients toward zero and risking underfitting of biologically meaningful signals. We therefore proceeded with a standard alpha = 1 for downstream comparisons, prioritizing interpretability of individual coefficients over the marginal gain in cross-validated R^2^.

With this setting, Ridge regression achieved R^2^ and MAE values comparable to the Random Forest model across split strategies (Figure S11B). However, direct comparison of average absolute error across all splitting strategies showed that Random Forest consistently outperformed Ridge regression, including spatially-aware, neighbor-excluded splits designed to test generalization across ocean regions (Figure S11C); Random Forest also showed lower maximum error across splits. Examination of the top Ridge coefficients (Figure S11D) further revealed that, aside from mOTUv4.0_002202 (the top sentinel *Pelagibacter* species identified in the main Random Forest analysis), the individual mOTU with the largest Ridge coefficients did not correspond to the most predictive taxa identified by Random Forest. Moreover, their raw abundance-temperature trajectories were considerably noisier (Figure S11E), which is consistent with Ridge regression distributing predictive weight across many weakly-informative features rather than isolating clear, biologically interpretable thermal signals.

In conclusion, Ridge regression achieved comparable overall predictive performance across split strategies, and coefficient exploration revealed a dilution of the biological signal across many features with weak individual effects, obscuring clear temperature-driven profiles. The RF model, therefore, proved superior not only in achieving lower maximum errors and better generalization in neighbor-excluded approaches, but also in effectively capturing and isolating ecologically interpretable thermal signatures of the marine microbiome.

#### Predictive performance of minimal feature sets across spatial splitting strategies

Reduced-feature models trained on only the four most explanatory mOTUs from the Random Forest approach were evaluated across all splitting strategies for both Random Forest and Ridge regression (Figure S12). Under 10-fold and 25-fold cross-validation, both models retained comparatively high predictive accuracy (R^2^ ≈ 0.81-0.82), consistent with the strong overall performance observed for the full-feature models. However, under spatially-aware splitting strategies (*i.e.*, including One-Dataset-Out, Longhurst, k-means, neighbor-excluded k-means, Ocean Region, and Ocean) R^2^ dropped sharply and consistently across both models (R^2^ ≈ 0.29-0.49), representing a substantial loss of explanatory power relative to the full-feature models. Notably, average absolute error remained comparatively stable and moderate across all splitting strategies (∼2.3-3.7 °C), including the space-aware ones, indicating that while a four-mOTU model can still produce numerically reasonable point predictions, it captures a substantially smaller share of the true variance in temperature once spatial structure is accounted for. These results reinforce the finding from the iterative drop-out analysis (Figure S9 and Figure S10) that robust generalization across ocean basins depends on a broader ensemble of predictive taxa, rather than a small, fixed set of top-ranked mOTUs.

### Supplementary Figures

**Fig S1.**
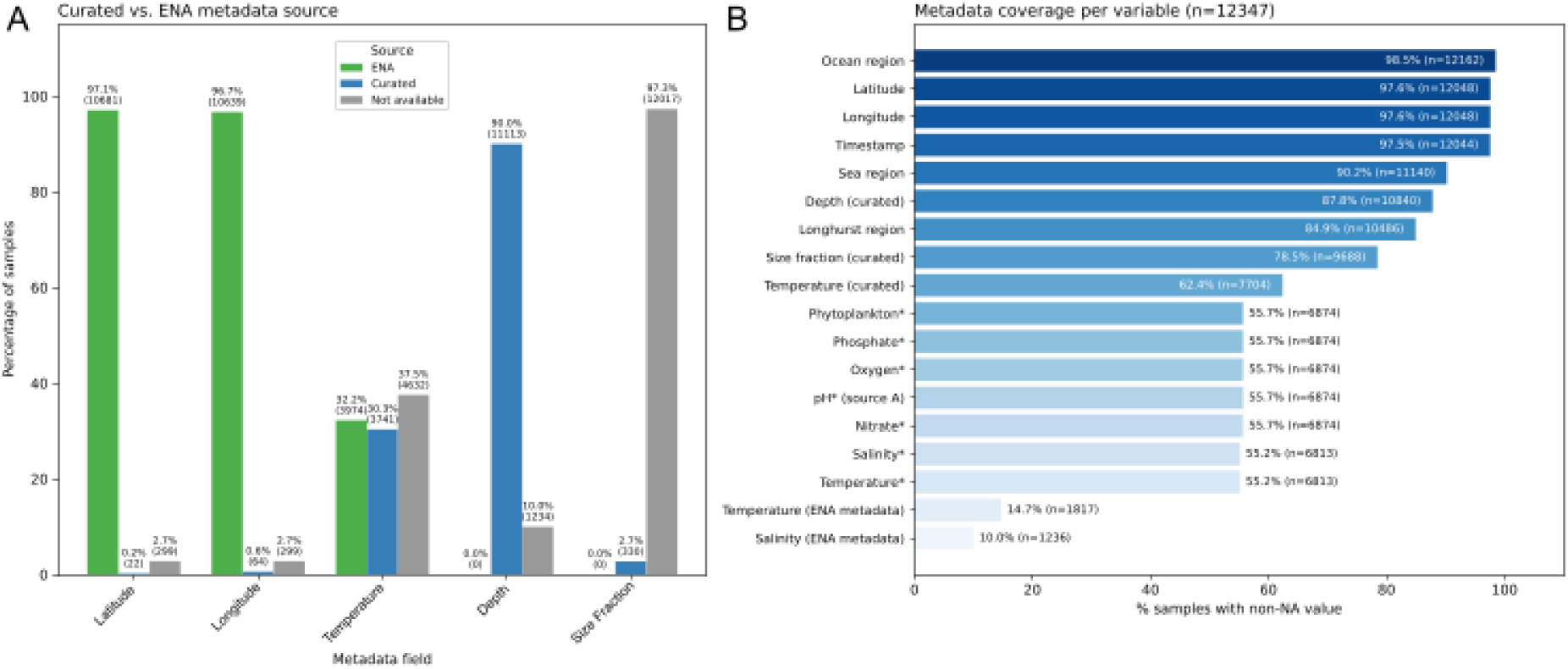
Metadata curation and collection. **(A)** Comparison of curated versus European Nucleotide Archive (ENA)-derived metadata for five key sample-level fields (latitude, longitude, temperature, depth, and size fraction; n = 12,347 total samples). For each field, bars show the percentage of samples where the curated value matched the ENA-reported value (“ENA”, green), where curation introduced a value differing from or absent in ENA (“Curated”, blue), and where no value was available from either source (“Not available”, gray). Latitude and longitude were rounded to the nearest integer degree prior to comparison to account for coordinate-precision differences between sources. Note depth and size fraction show a disproportionately large “Curated” fraction relative to temperature and coordinates; this reflects unit and format harmonization (*e.g.*, standardizing depth to meters and consolidating inconsistent size-fraction notations across studies) rather than genuine disagreement in the underlying measured values. **(B)** Overall metadata coverage (percentage of non-missing values) across all fields used in downstream analyses, including manually curated in-situ measurements, ENA-derived metadata, and satellite-derived environmental variables from Copernicus Marine Service (marked with *). Bar labels indicate percentage and absolute sample count (n) for each field.

**Fig S2.**
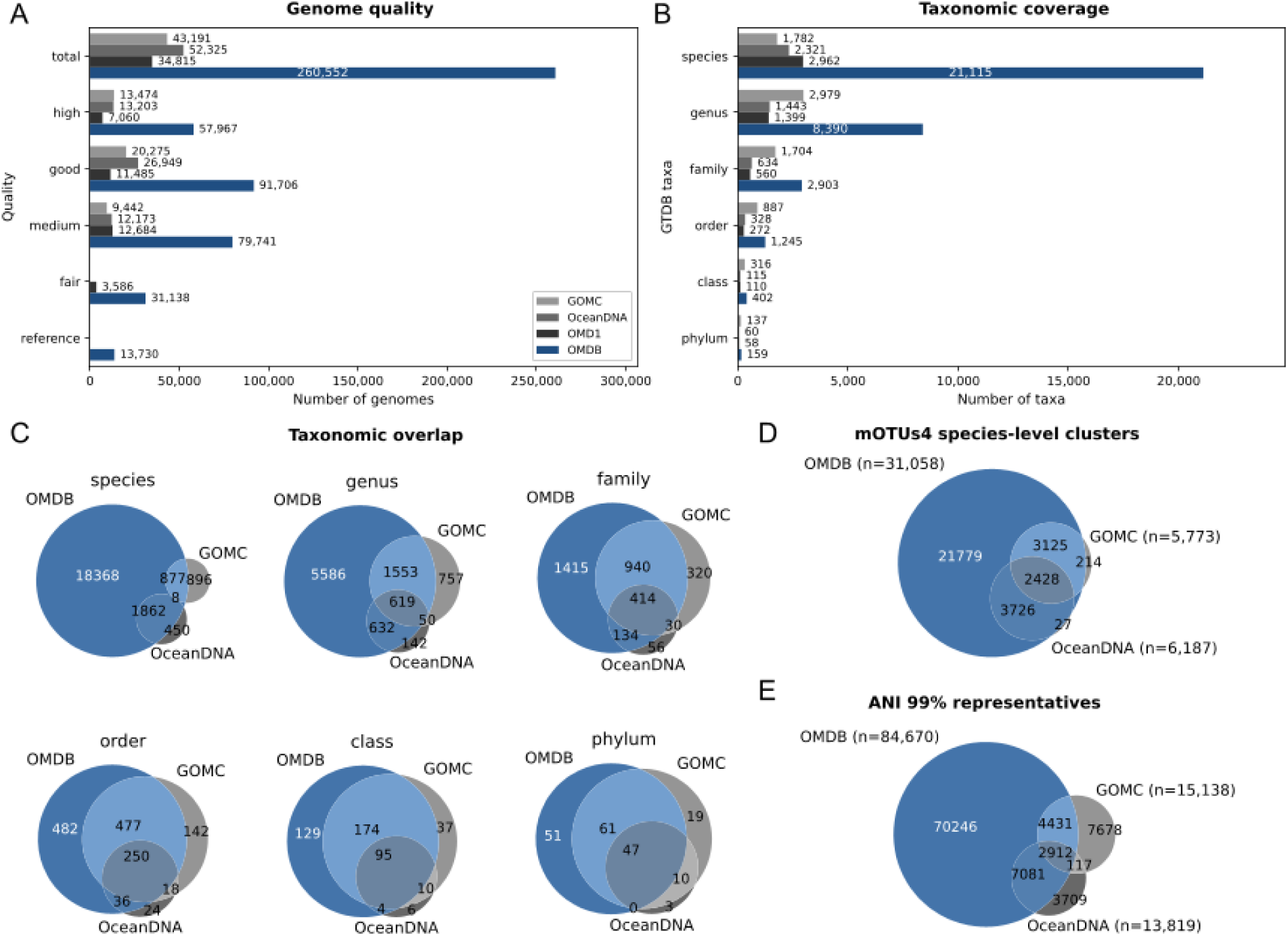
OMDB-GOMC-OceanDNA resources comparative. **(A)** Number of genomes by CheckM2 quality tier (reference, fair, medium, good, high, total) and **(B)** number of GTDB taxa represented at each taxonomic rank, compared across OMDB, OMD1, OceanDNA, and GOMC. **(C)** Venn diagrams showing overlap in taxa identified by each database at the species, genus, family, order, class, and phylum level. **(D)** Overlap in mOTUs4 species-level clusters for the genomes that were assigned to an existing mOTU, and **(E)** dereplicated genome representatives at 99% average nucleotide identity (ANI) across the three databases.

**Fig S3.**
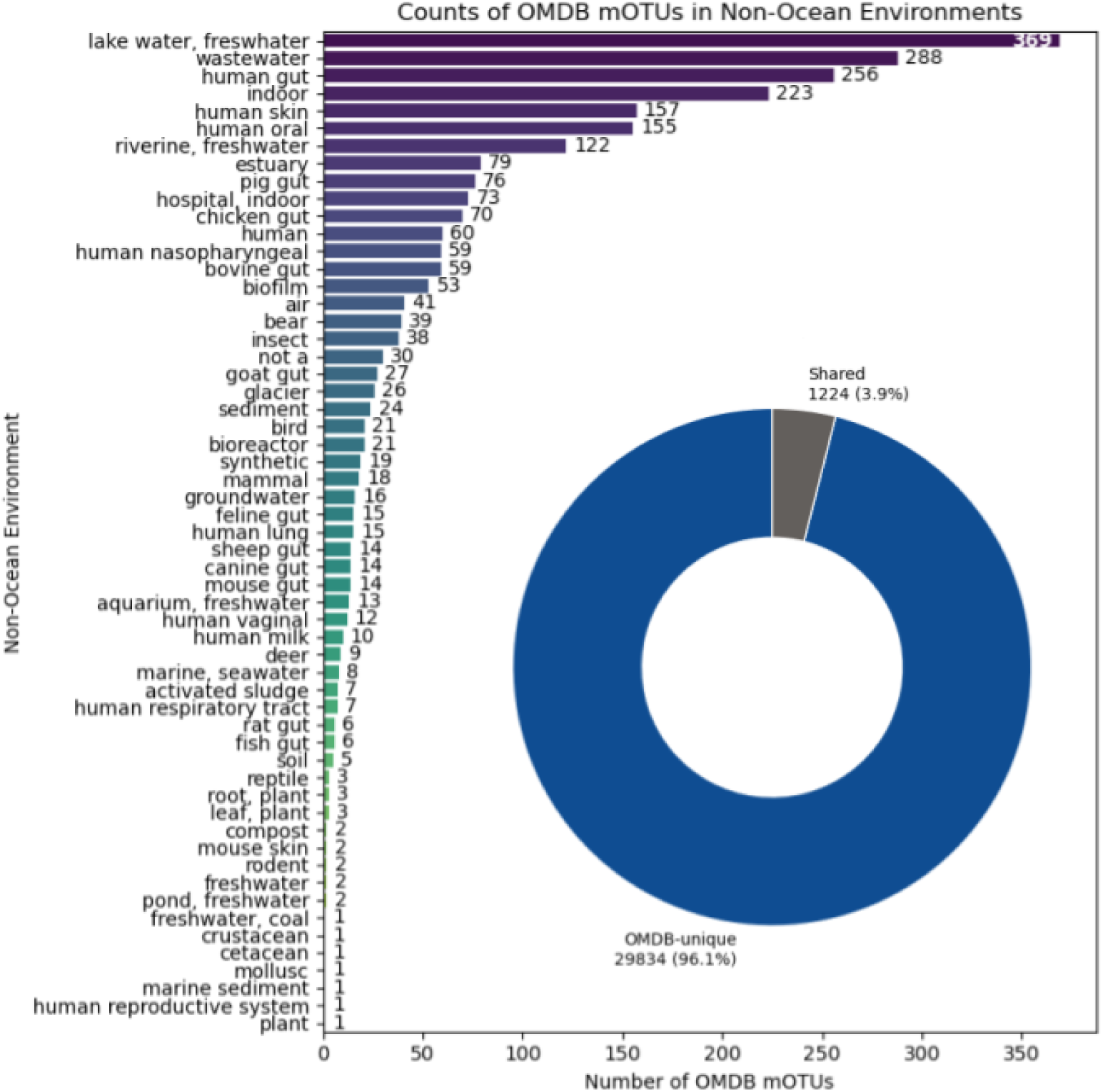
Extension to other environments Distribution of OMDB mOTUs across non-ocean environments. Horizontal bar chart showing the number of OMDB mOTUs also detected in non-ocean environments ranked in descending order. The inset donut chart summarizes the mOTUs overlap between ocean and non-ocean environments, showing mOTUs unique to OMDB/ocean samples (dark blue, 29,834; 96.1%) and mOTUs shared with at least one non-ocean environment (grey, 1,224; 3.9%).

**Fig S4.**
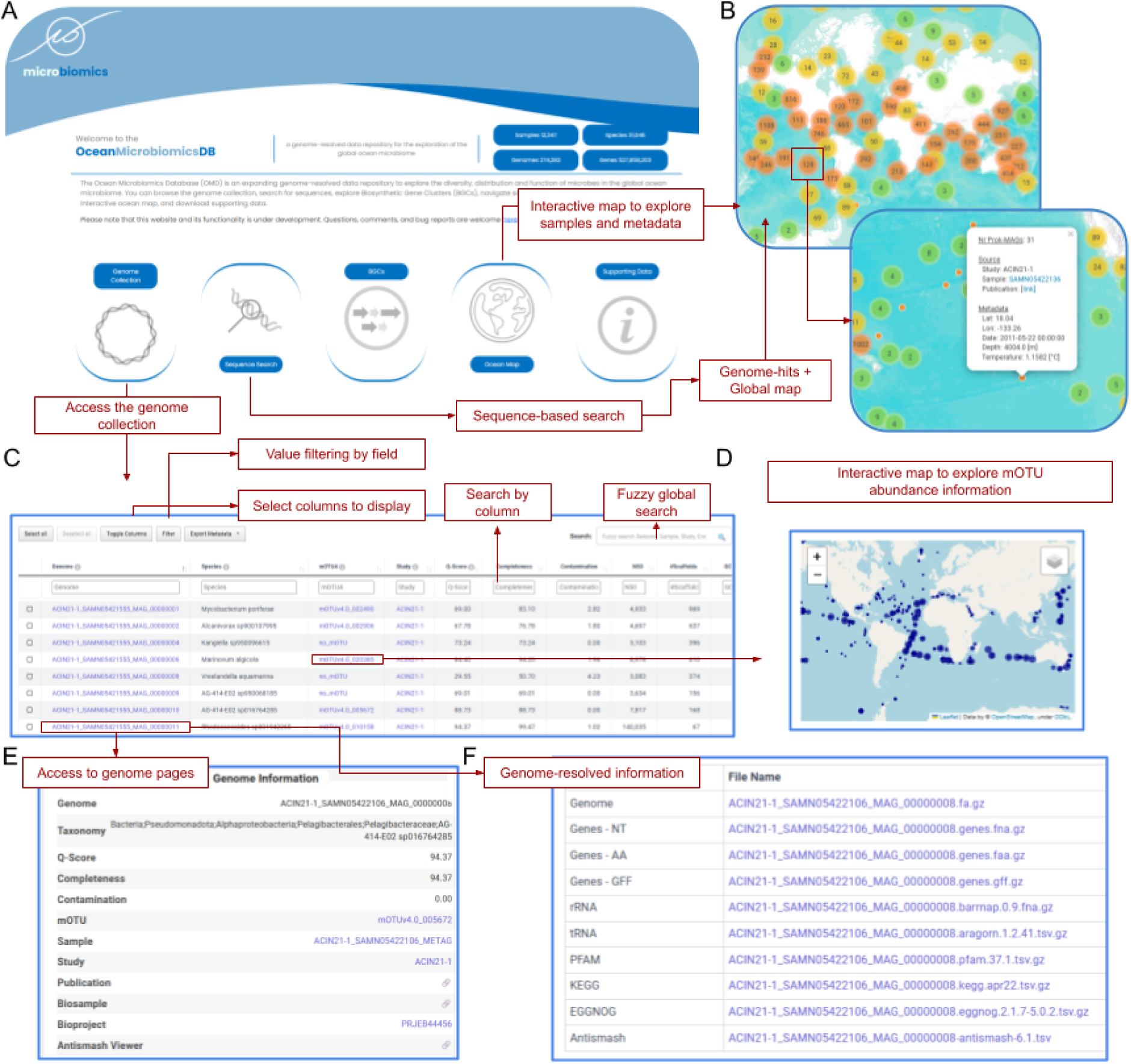
Overview of the OMDB web interface. **(A)** Landing page providing access to the genome collection and summary statistics (samples, species, genomes, and genes), together with navigation to the main tools: Genome Collection, Sequence Search, BGCs, Ocean Map, and Supporting Data. **(B)** Map-based search displaying samples, variables and linking to their genome on an interactive world map, enabling exploration of sample distribution and associated metadata. **(C)** Genome collection browser supporting per-field value filtering, column selection/toggling, per-column search, and fuzzy global search across genome-level metrics. These ‘collection’ pages are also available per mOTUs, sample and study. **(D)** Interactive map for exploring per-mOTU abundance and geographic distribution. **(E)** Individual genome page example, displaying diverse information, metrics, and links to the associated sample, study, and publication **(F)** Genome-resolved data downloads providing direct access to genome sequences, predicted genes (nucleotide and amino acid), GFF annotations, functional annotation outputs (rRNA, tRNA, PFAM, KEGG, EGGNOG), and antiSMASH results for each genome.

**Fig S5.**
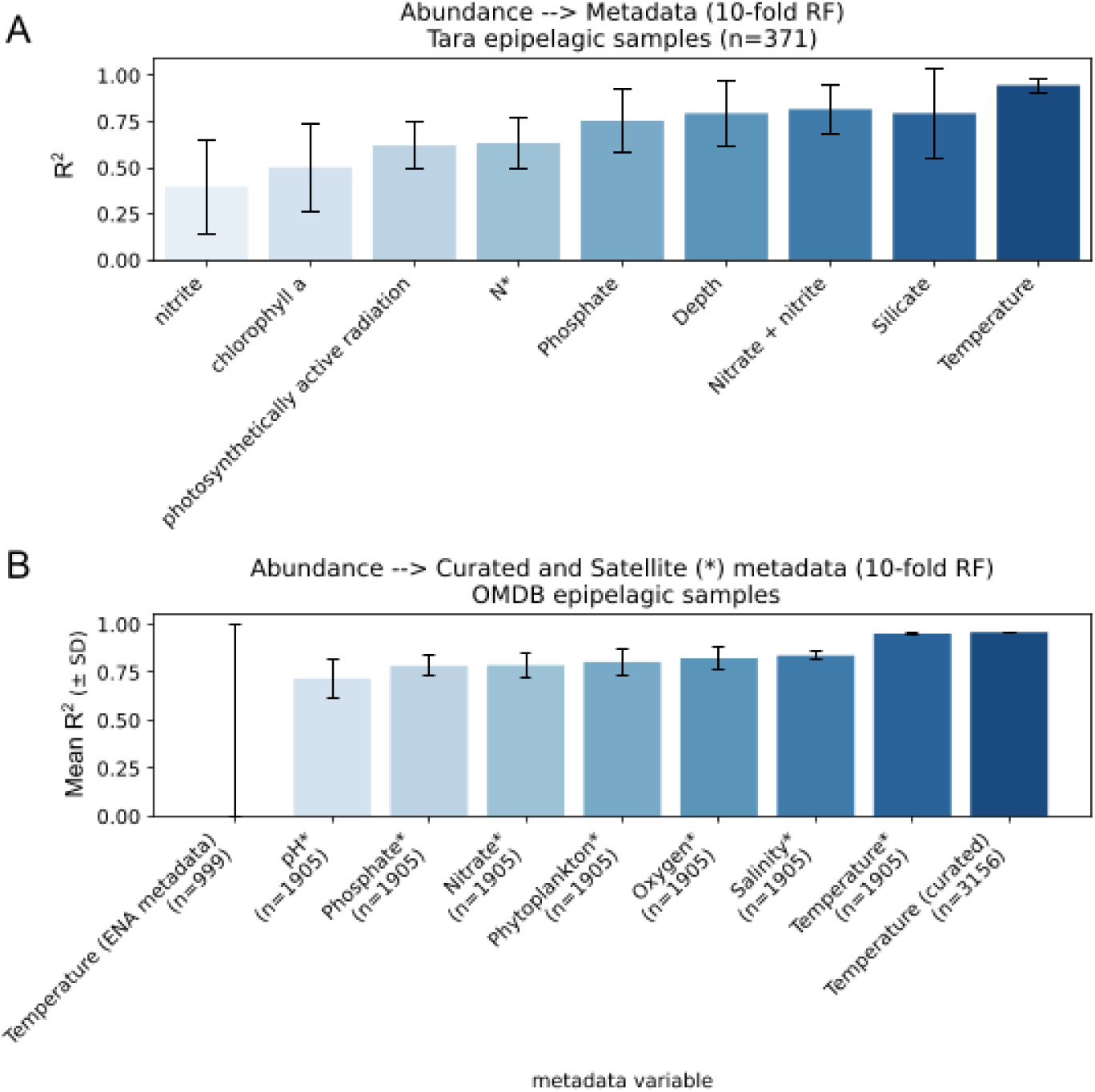
*Tara* and satellite-derived metadata predictability from microbial abundance data. Mean R^2^ (± SD) across 10-fold cross-validation Random Forest models predicting each environmental variable from community abundance data. **(A)** *Tara* Oceans epipelagic samples (n = 371), using in-situ measured metadata. **(B)** OMDB epipelagic samples, using curated in-situ temperature and satellite-derived (Copernicus Marine Service) environmental variables (marked with *), compared against raw ENA metadata-derived temperature annotations; sample sizes (n) for each variable are indicated below the corresponding bar.

**Fig S6.**
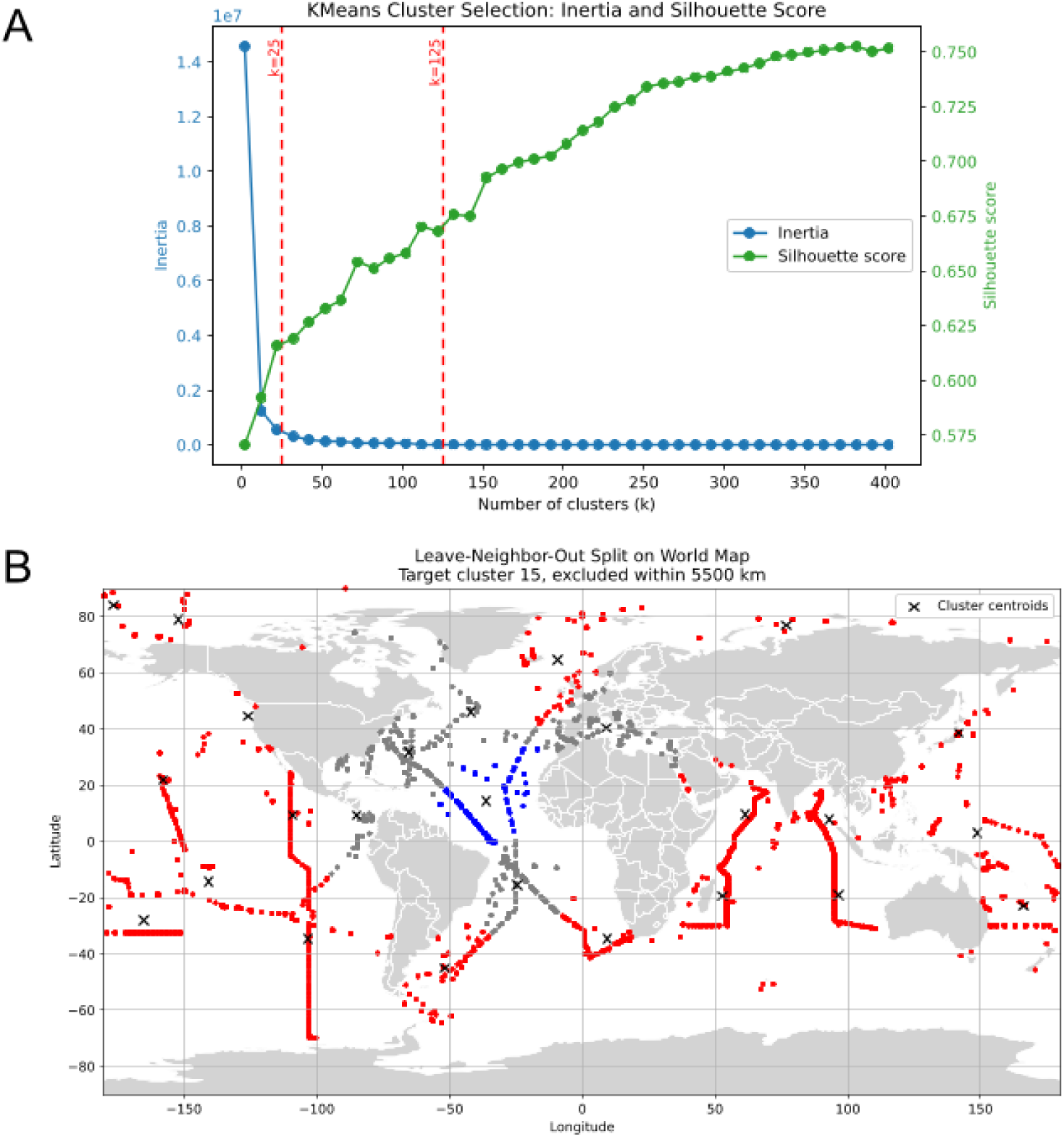
Spatial splitting strategy for Leave-Neighbor-Out (LNO) cross-validation. **(A)** K-means clustering was performed directly on sample latitude/longitude coordinates across a range of k values (k = 2-410, in increments of 10) and evaluated using two complementary metrics: inertia (within-cluster sum of squared distances; blue, left axis) and silhouette score (green, right axis). Inertia decreases monotonically as k increases, whereas the silhouette score increases steadily across the range examined. Dashed red lines indicate the two k values selected for downstream spatial cross-validation analyses (k = 25 and k = 125). **(B)** Example map illustrating the LNO spatial cross-validation scheme with k = 25. Samples were grouped into k-means clusters based on geographic coordinates, with each cluster sequentially designated as a held-out test set. For each target cluster (test set, blue), training samples (red) within a fixed exclusion radius of the cluster centroid were removed to reduce spatial autocorrelation and information leakage. Black crosses mark cluster centroids. For cluster 15, the 5,500 km radius corresponds to the minimum distance between neighboring centroids, ensuring a consistent spatial buffer. This procedure was repeated for both clustering resolutions (k = 25 and k = 125), with each cluster held out once for testing.

**Fig S7.**
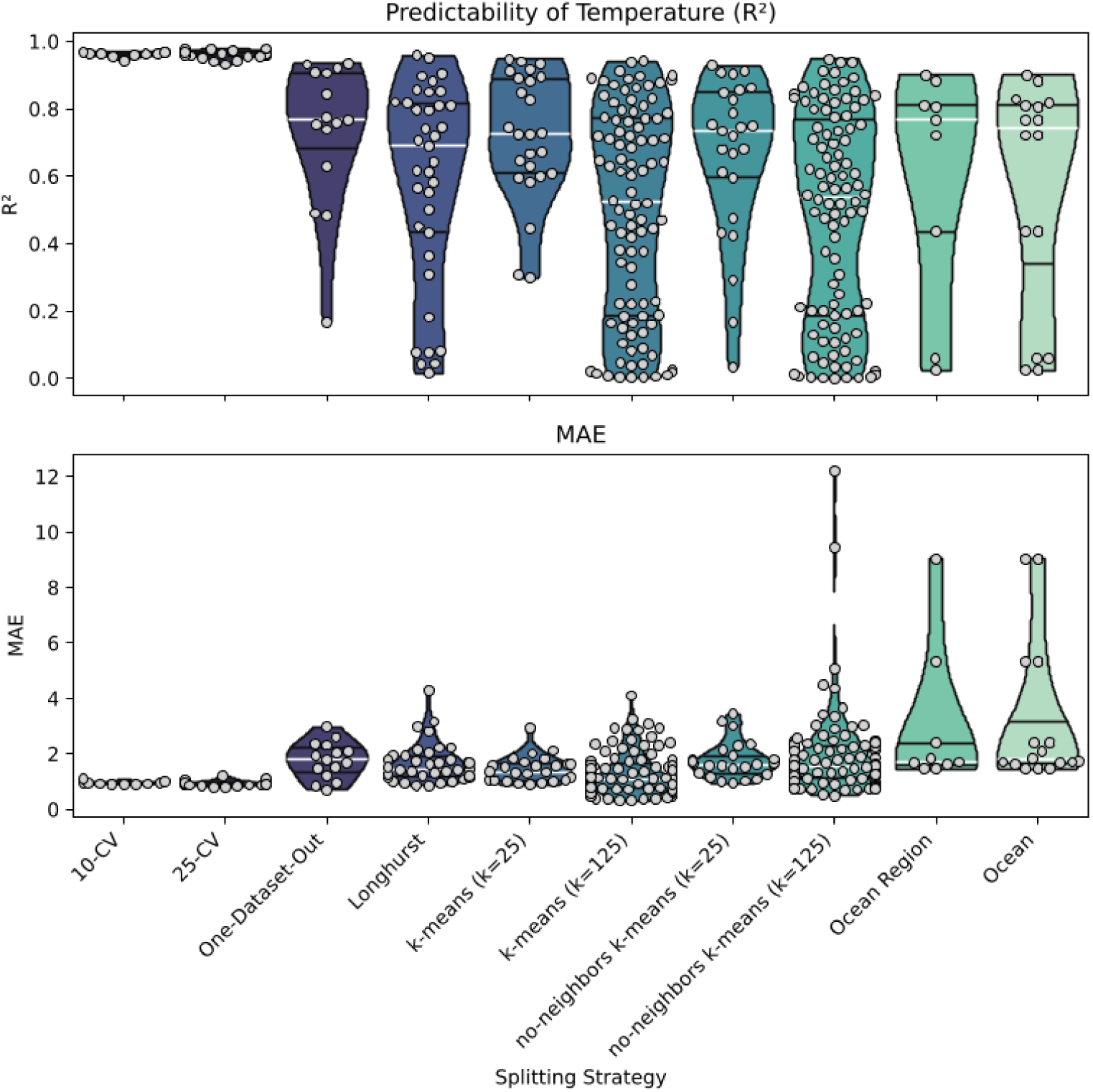
Predictability of temperature under increasingly spatially independent train/test splitting strategies. Random Forest performance for predicting temperature from community abundance data, evaluated using 10-fold repeated testing under ten train/test splitting strategies that progressively increase spatial separation between training and test sets. **(Top)** R^2^ correlation coefficient for each test fold or held-out group. **(Bottom)** Mean absolute error (MAE, °C) for each test fold or held-out group. Violin plots show the distribution of values across folds or groups for each strategy; white lines indicate medians, while black lines indicate the interquartile range. Individual grey points represent the result for each fold or held-out group,

**Fig S8.**
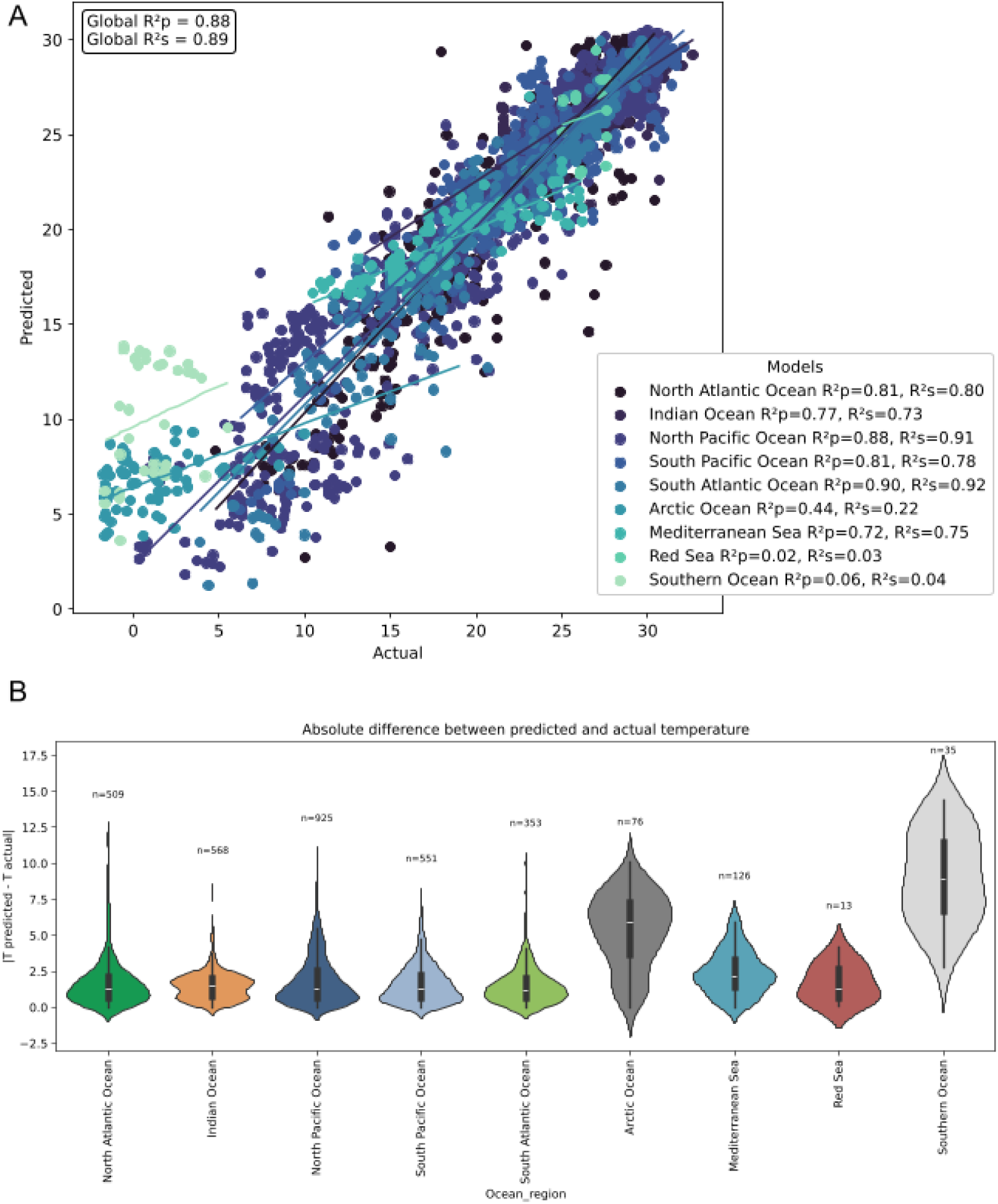
Predicted versus actual temperature by ocean region. **(A)** Scatter plot of predicted versus actual temperature (°C) across all samples, color-coded by ocean region with per-region linear regression fits. Global Pearson (R^2^p) and Spearman (R^2^s) correlations are displayed in the top-left inset, with per-region metrics provided in the legend. Accuracy is strong for the major ocean basins (North Atlantic, Indian, North Pacific, South Pacific, South Atlantic and the Mediterranean Sea, but markedly lower for the Arctic Ocean and Southern Ocean. The Red Sea R^2^ values are low due the low number of samples (n = 13). **(B)** Violin plots showing absolute prediction error (°C) by ocean region, with sample sizes labeled above each violin. Mirroring panel A, prediction errors are small and tightly clustered for major basins, moderate for the Mediterranean Sea, and substantially higher and more variable for the Arctic and Southern Oceans.

**Fig S9.**
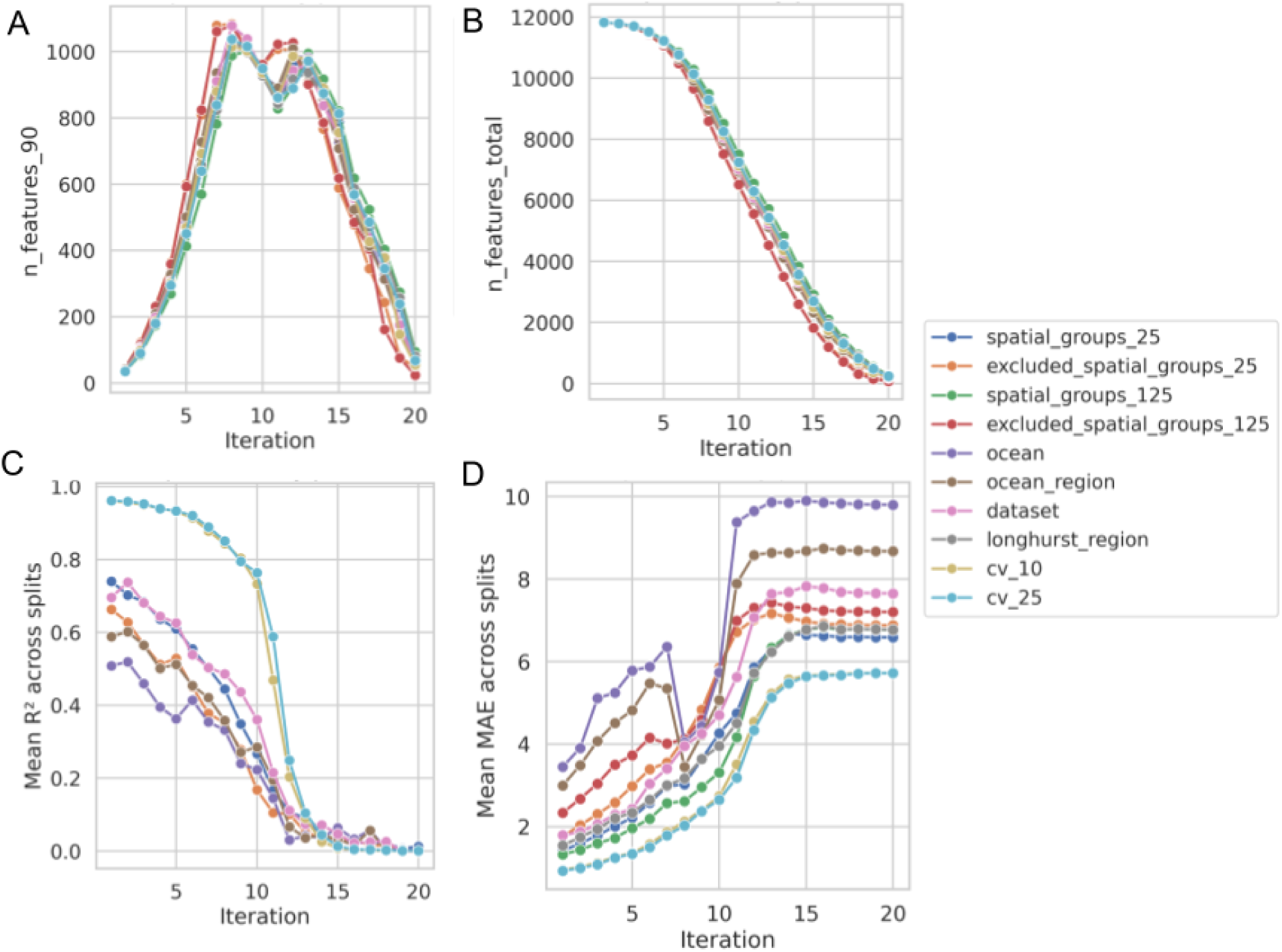
Iterative drop-out Random Forest analysis. Iterative feature drop-out analysis evaluating Random Forest predictive redundancy. At each iteration N+1, mOTUs accounting for 90% of the cumulative feature importance at iteration N were removed before retraining on the remaining features. This process was repeated across 20 iterations for each splitting strategy. **(A)** Number of mOTUs required to reach the 90% threshold at each iteration. **(B)** Total mOTUs remaining after cumulative drop-outs at each iteration. **(C)** Mean Pearson R^2^ across cross-validation splits at each iteration. **(D)** Mean absolute error (MAE, °C) across cross-validation splits at each iteration.

**Fig S10.**
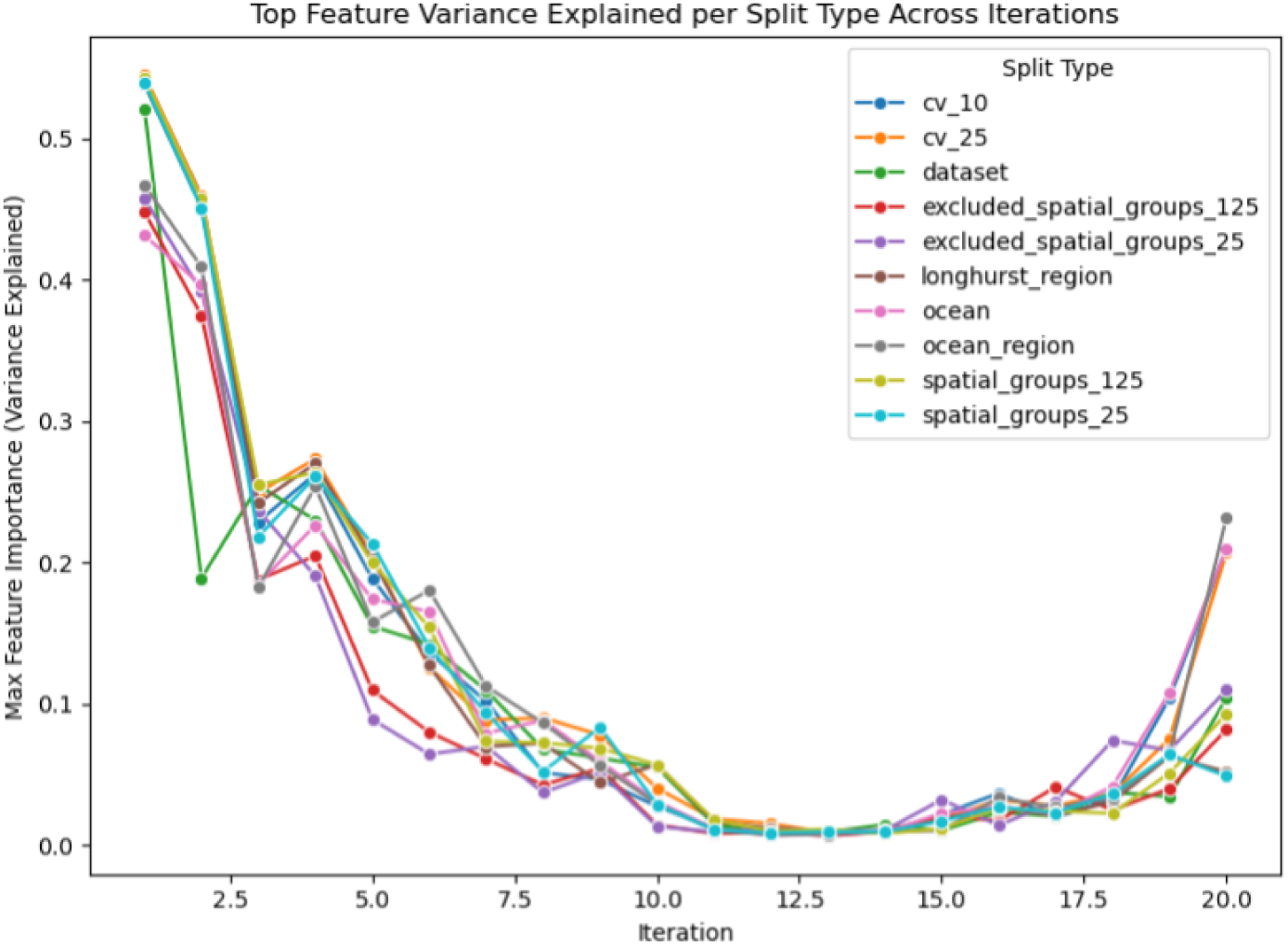
Predictive contribution of the single most important mOTU across drop-out iterations. For each iteration of the feature drop-out analysis described in Figure S9, the variance explained by the single top-ranked mOTU (maximum individual feature importance) is shown across all splitting strategies.

**Fig S11.**
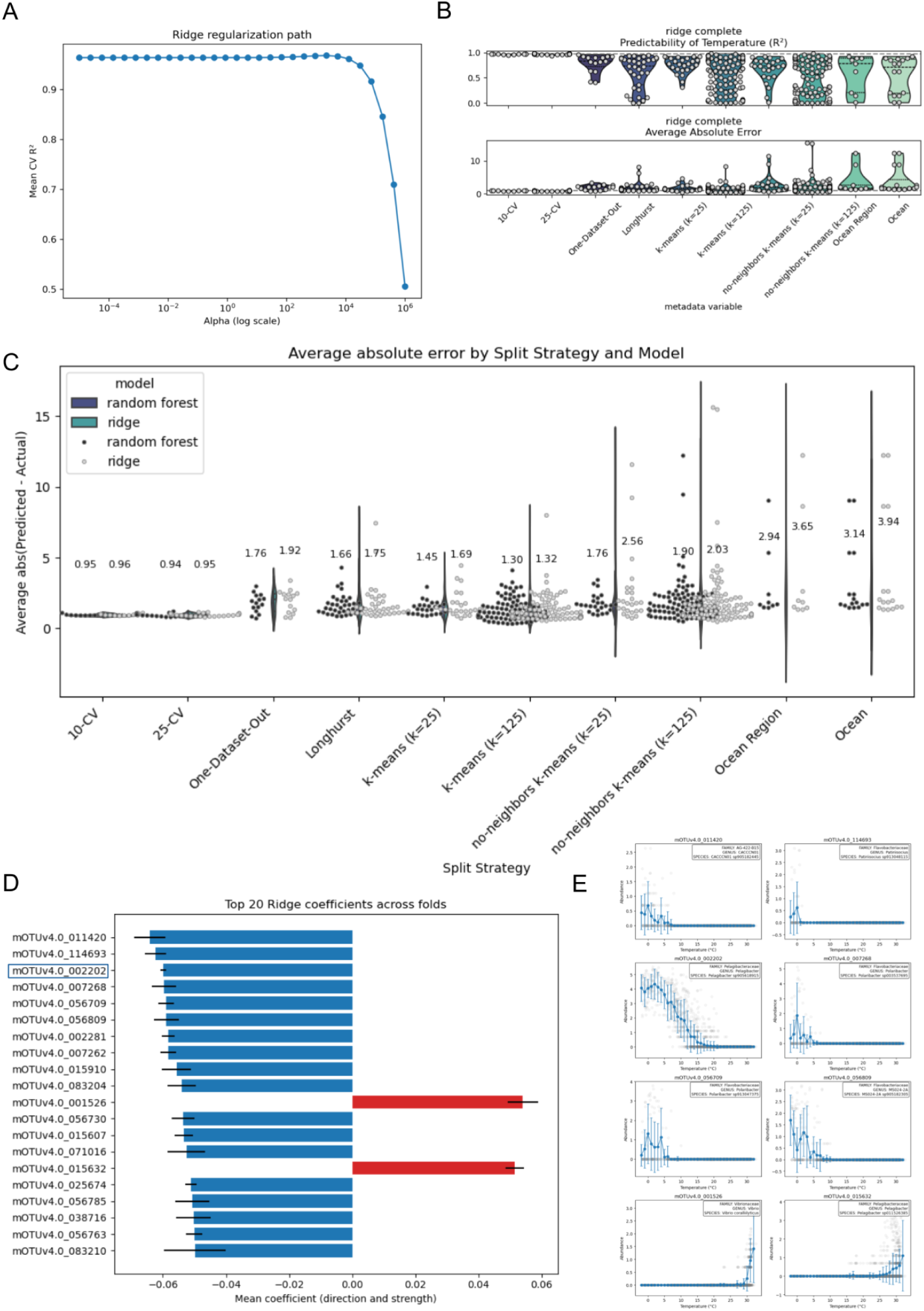
Benchmarking of Ridge regression against Random Forest for temperature prediction. **(A)** Ridge regularization path evaluated using 10-fold cross-validation, showing mean cross-validated R^2^ across values of the regularization parameter alpha (log scale). **(B)** Temperature predictability (R^2^, top) and mean absolute error (bottom) for the Ridge regressor (alpha = 1) across splitting strategies, evaluated using the same framework as the Random Forest model. **(C)** Distribution of absolute prediction error (|predicted - actual|, °C) across split strategies for Random Forest (dark violins) and Ridge (light violins/points), with mean absolute error indicated above each distribution. **(D)** The 20 largest Ridge regression coefficients (mean ± SD across folds), ranked by absolute magnitude. The two largest coefficients, mOTUv4.0_001526 and mOTUv4.0_071016 (highlighted in red), were substantially larger than the remaining coefficients. **(E)** Relationship between raw abundance and temperature for eight of the mOTUs receiving the largest Ridge weights. With the exception of mOTUv4.0_002202, these relationships are visually noisy and do not exhibit the sharply constrained thermal peaks characteristic of the top-ranked Random Forest taxa (Figure 2E).

**Fig S12.**
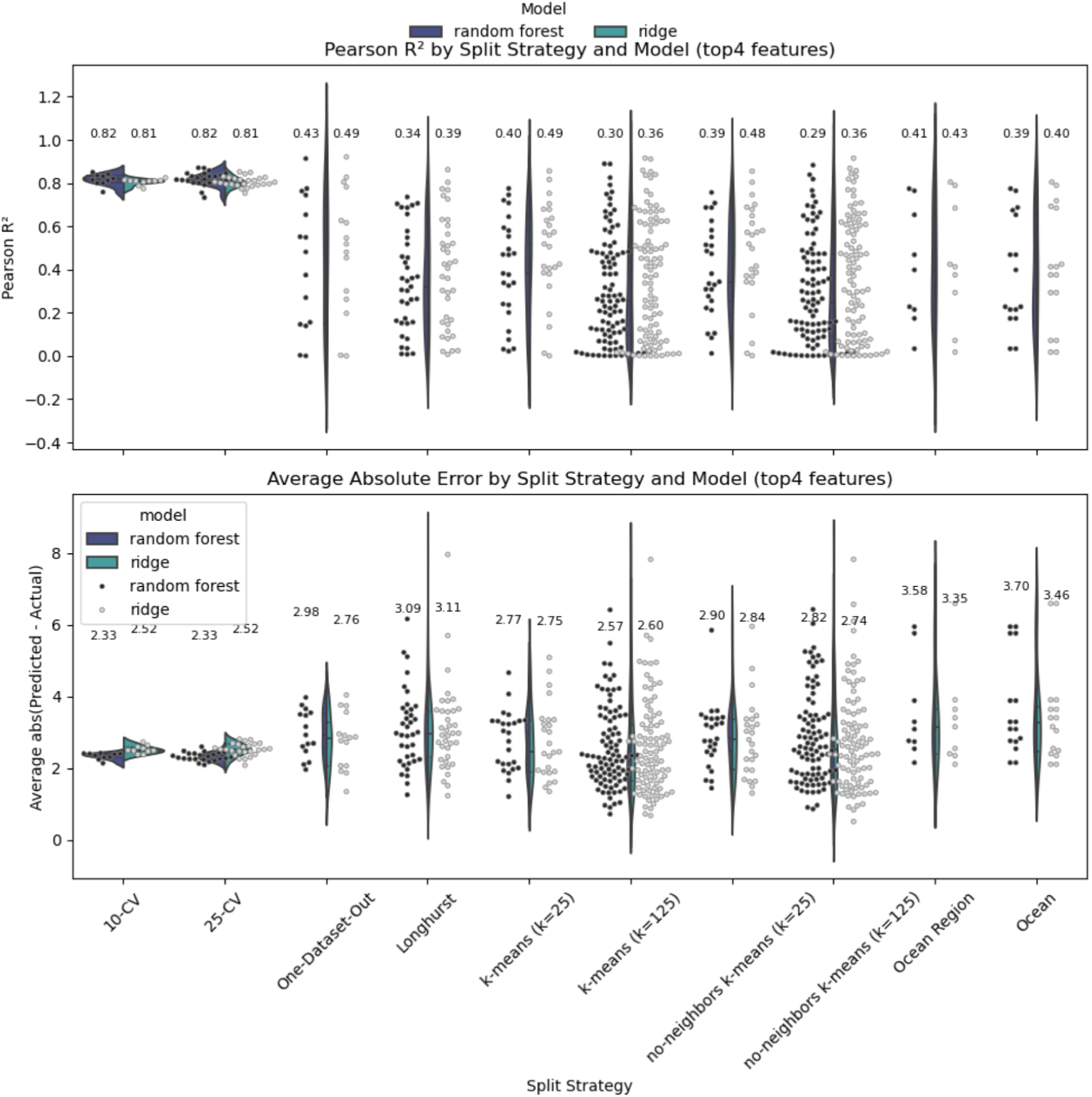
Reduced-feature model performance using the four most explanatory mOTUs. Random Forest and Ridge regression models were retrained using exclusively the top four mOTUs by feature importance, and evaluated across all splitting strategies. **(Top)** Pearson R^2^ by split strategy and model; mean values are annotated above each distribution.. **(Bottom)** MAE (°C) by split strategy and model; mean values are annotated above each distribution. Together, these results indicate that while a minimal four-mOTU model can still achieve tolerable average error, its ability to explain variance collapses under spatially structured validation, demonstrating that a broader ensemble of predictive mOTUs (beyond this minimal top-ranked set) is essential to maintain robust predictability across ocean basins.

**Fig S13.**
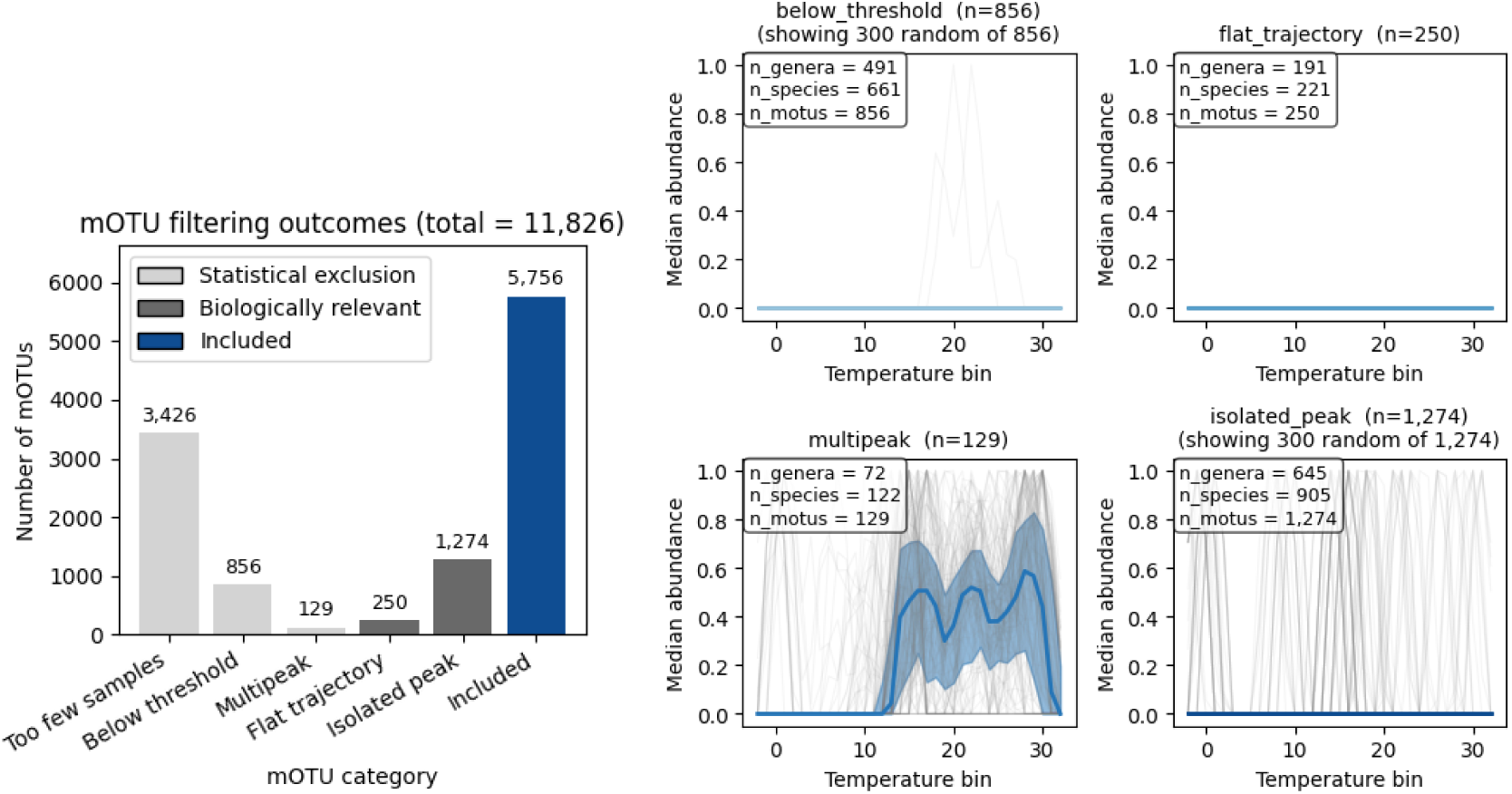
Trajectories of mOTUs excluded from thermal shape-clustering, categorized by exclusion reason. (**Left**) Number of mOTUs assigned to each exclusion category during quality filtering of abundance-temperature trajectories. (**Right**) Panels display smoothed, min-max normalized abundance trajectories across 1 °C temperature bins for individual mOTUs (light gray lines), alongside category medians (solid lines) and interquartile ranges (shaded bands, 25th-75th percentiles). Panel titles state the exclusion category, total mOTUs (n), and the number of randomly subsampled trajectories plotted when n>300 (summary statistics and counts reflect all mOTUs). Inset boxes detail the number of distinct genera, species, and mOTUs per category. Criteria fall into two groups: statistical exclusions due to insufficient or ambiguous signal (below threshold, multipeak) and biologically informative exclusions lacking a single identifiable thermal optimum (isolated peak, monotonic cold, monotonic warm, flat trajectory).

**Fig S14.**
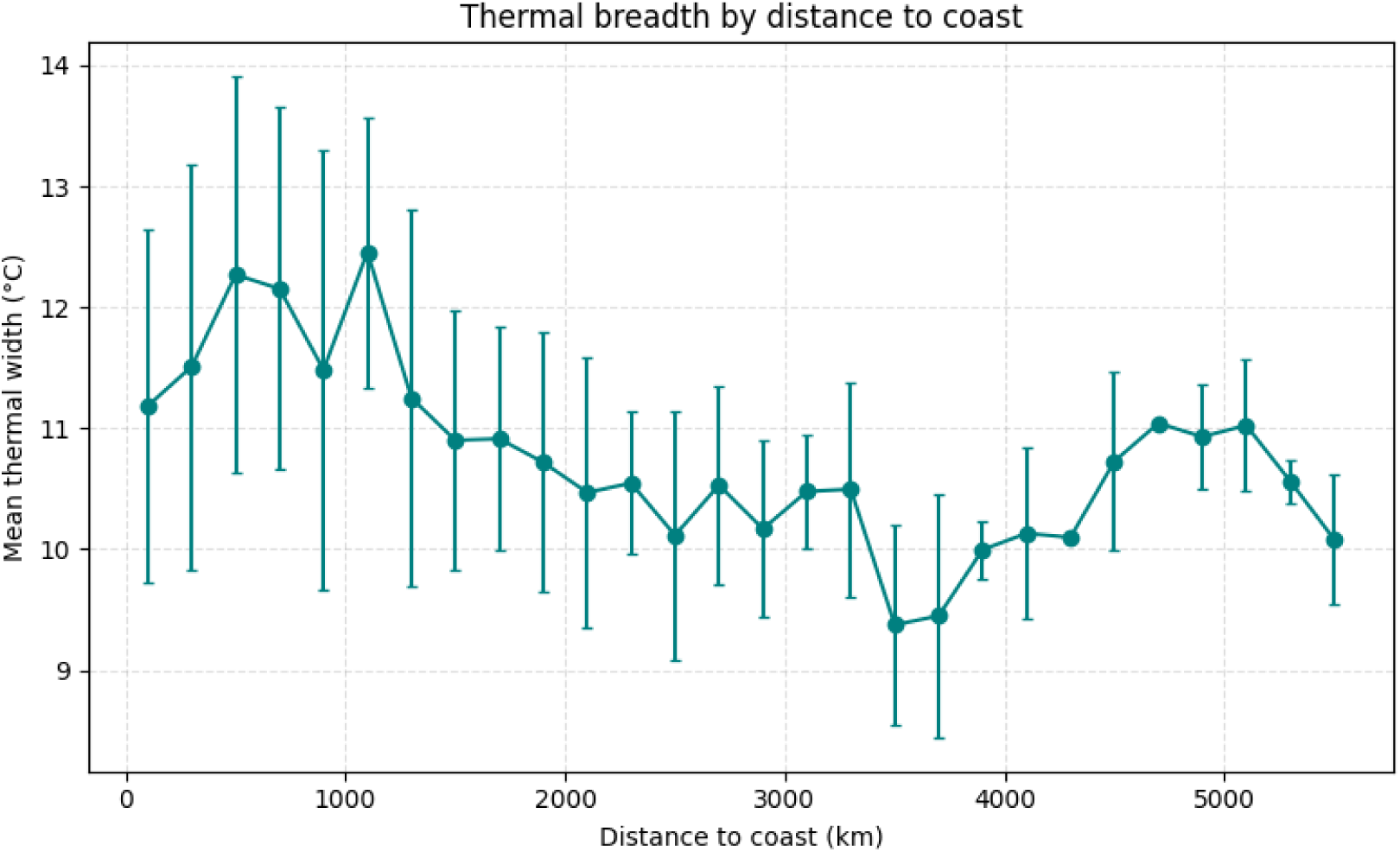
Coastal proximity is associated with broader thermal niches. Mean thermal niche breadth and standard deviation as error bars of ocean microbial mOTUs (Y-axis), binned by distance to the nearest coastline (km; X-axis).

**Fig S15.**
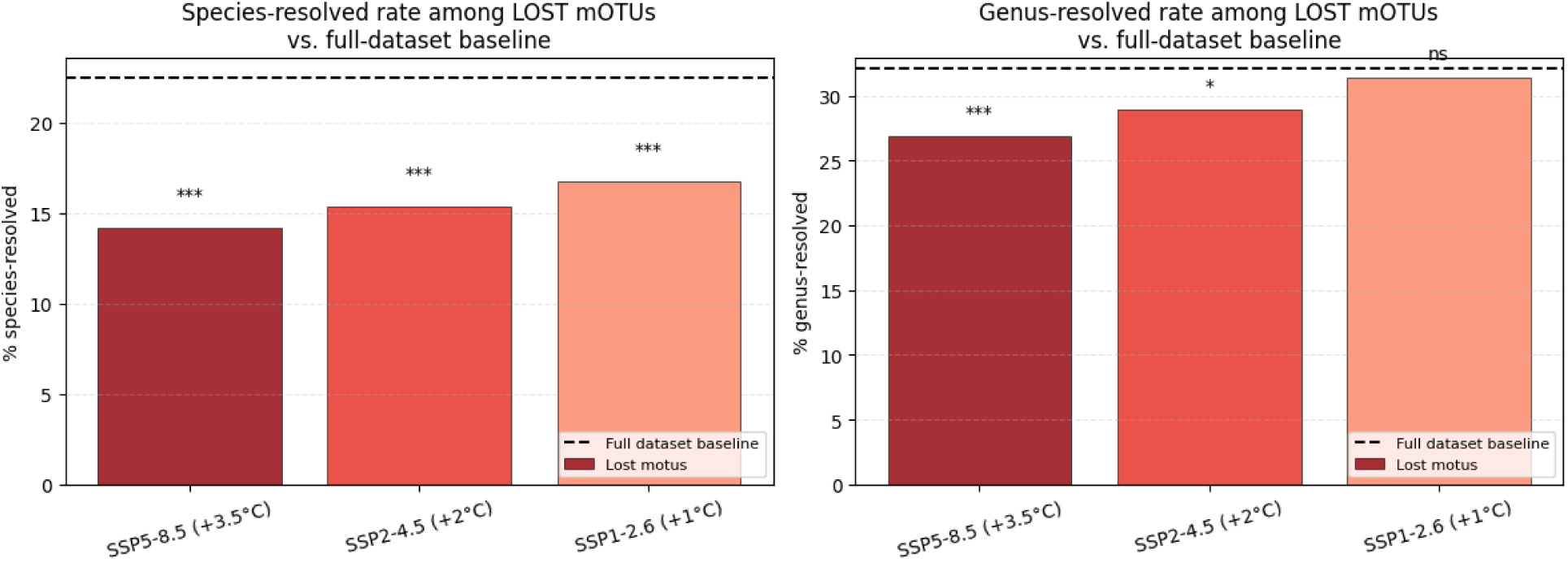
Taxa lost to warming are disproportionately taxonomically unresolved. Percentage of mOTUs carrying a formal species-level (left) or genus-level (right) classification among mOTUs projected to be lost under each IPCC warming scenario, compared to the full-dataset baseline (dashed line). Asterisks denote significance vs. baseline (Fisher’s exact test, Benjamini-Hochberg corrected; **p<0.01, ***p<0.001; ns, not significant).

**Fig S16.**
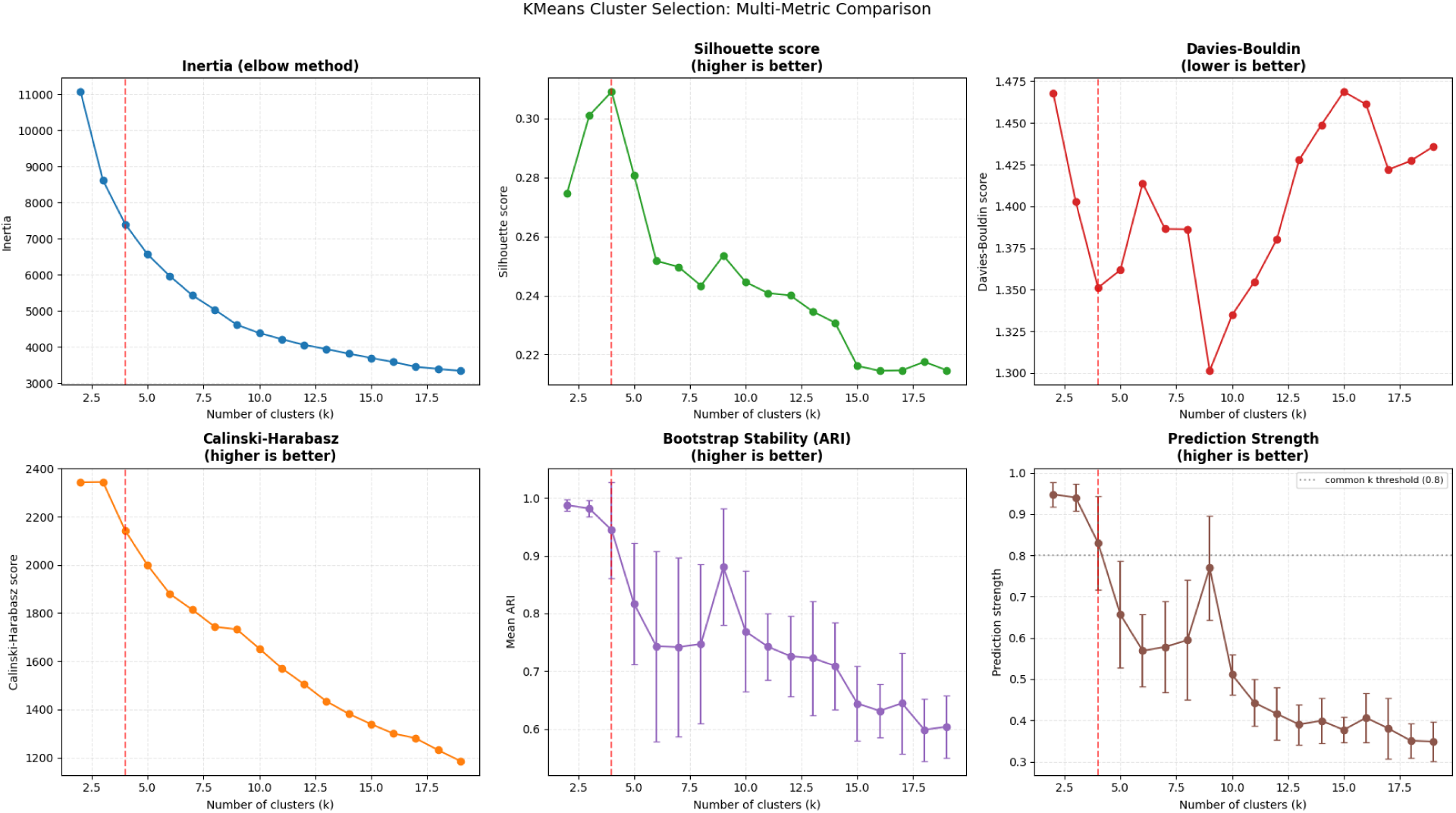
Selection of number of informative clusters based on mOTUs abundance response in a temperature gradient. Evaluation of K-means clustering solutions across k = 2-19 clusters, using six complementary metrics: inertia (within-cluster sum of squared distances; elbow method), silhouette score, Davies-Bouldin index, Calinski-Harabasz index, bootstrap stability (mean adjusted Rand index, ARI, across resampled datasets; error bars show standard deviation), and prediction strength (Tibshirani & Walther, 2005; error bars show standard deviation across repeated train/test splits). The dashed vertical line marks k = 4, the selected number of clusters.

**Fig S17.**
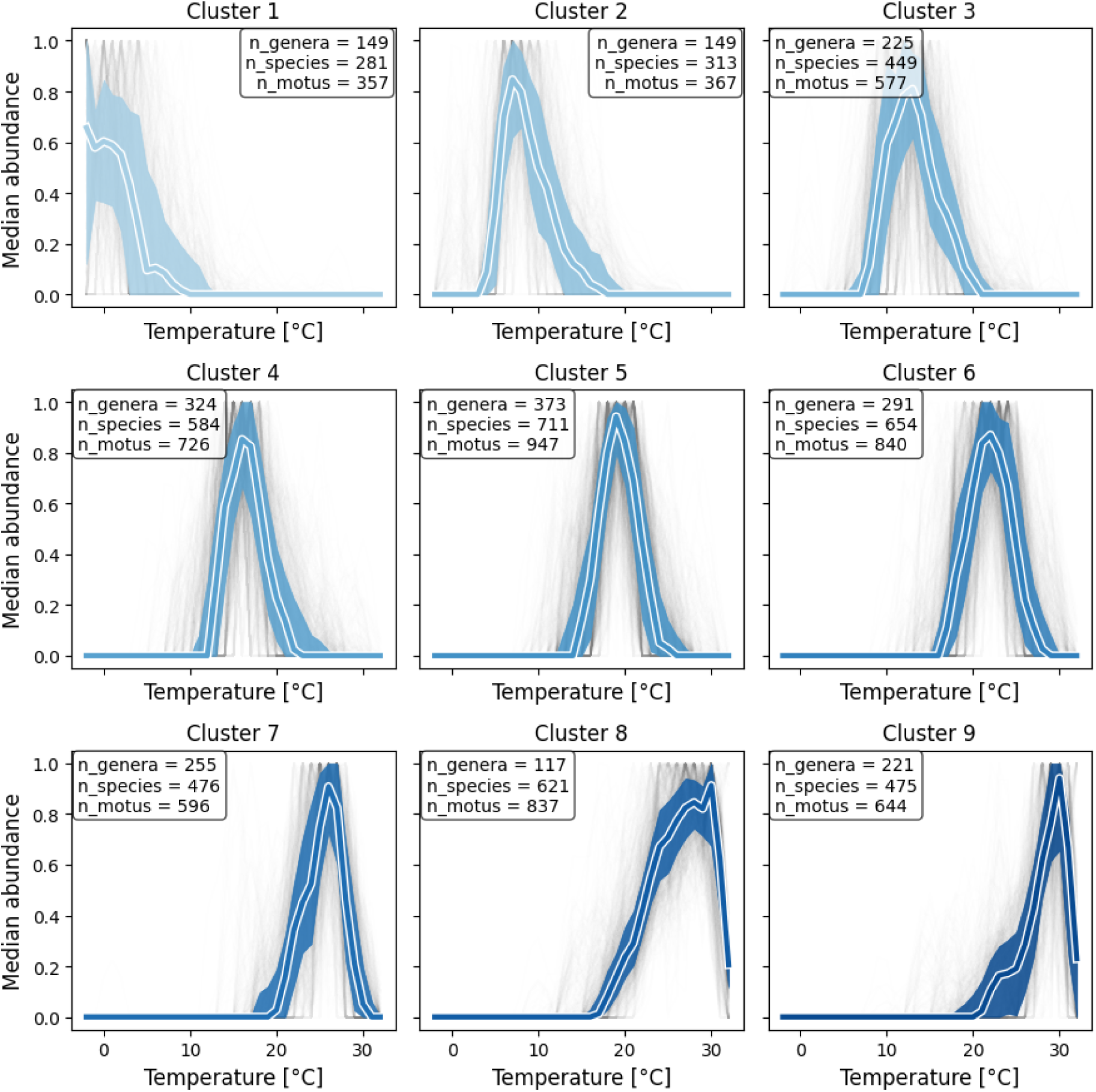
K-means clustering at k = 9 illustrates over-partitioning of the thermal gradient. Median (bold line) and interquartile range (shaded band) of normalized abundance trajectories for the nine clusters obtained under a k=9 solution, ordered by ascending peak temperature. Gray lines show individual mOTU trajectories within each cluster; inset boxes report the number of distinct genera, species, and mOTUs per cluster. Despite showing a secondary local maximum in prediction strength (Figure S16), the k = 9 solution does not resolve nine qualitatively distinct thermal response shapes.

**Fig S18.**
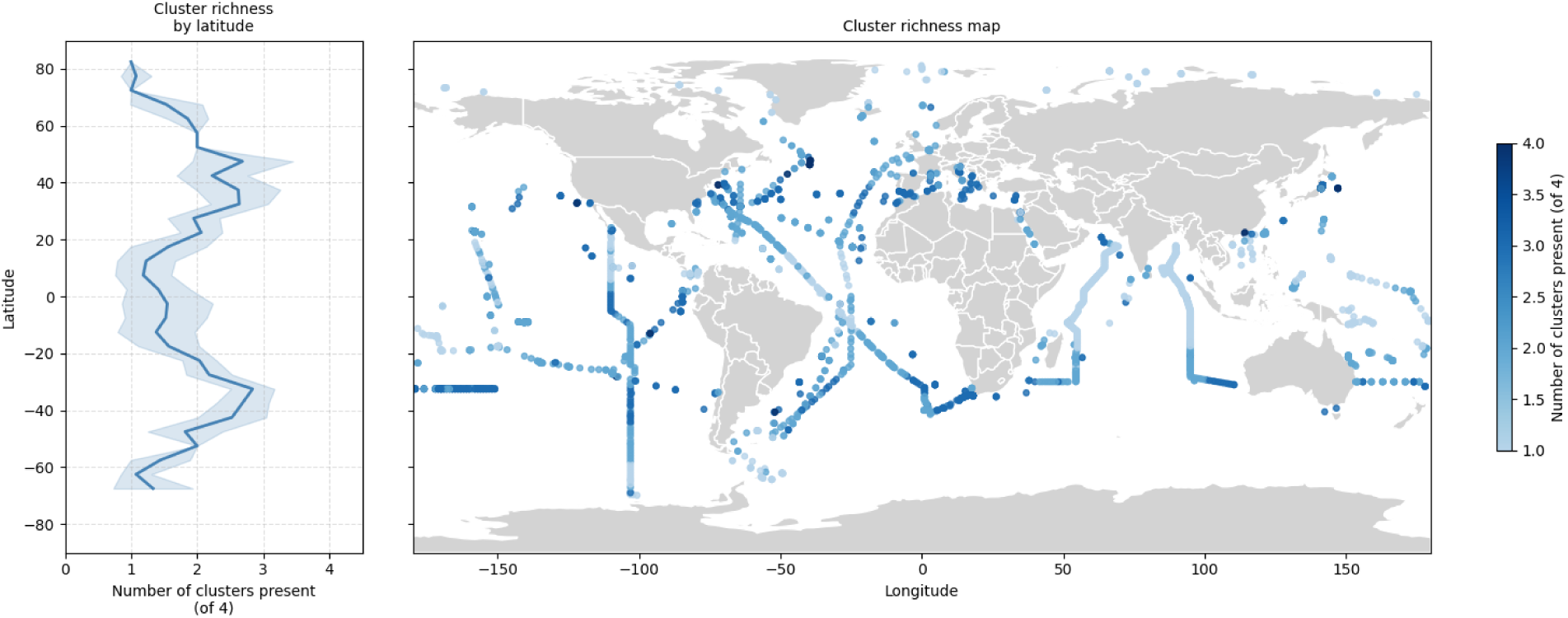
Local co-occurrence of thermal clusters across latitude and geography (Left) Mean number of distinct thermal clusters present per sample (out of 4 total; presence defined as a cluster contributing ≥5% of the local relative abundance) as a function of latitude, binned in 5° increments; the shaded band shows ±1 SD. **(Right)** Geographic distribution of the same metric for each individual sample. Color intensity indicates the number of co-occurring thermal clusters at that location, from 1 (single cluster dominates locally) to 4 (all clusters represented).

**Fig S19.**
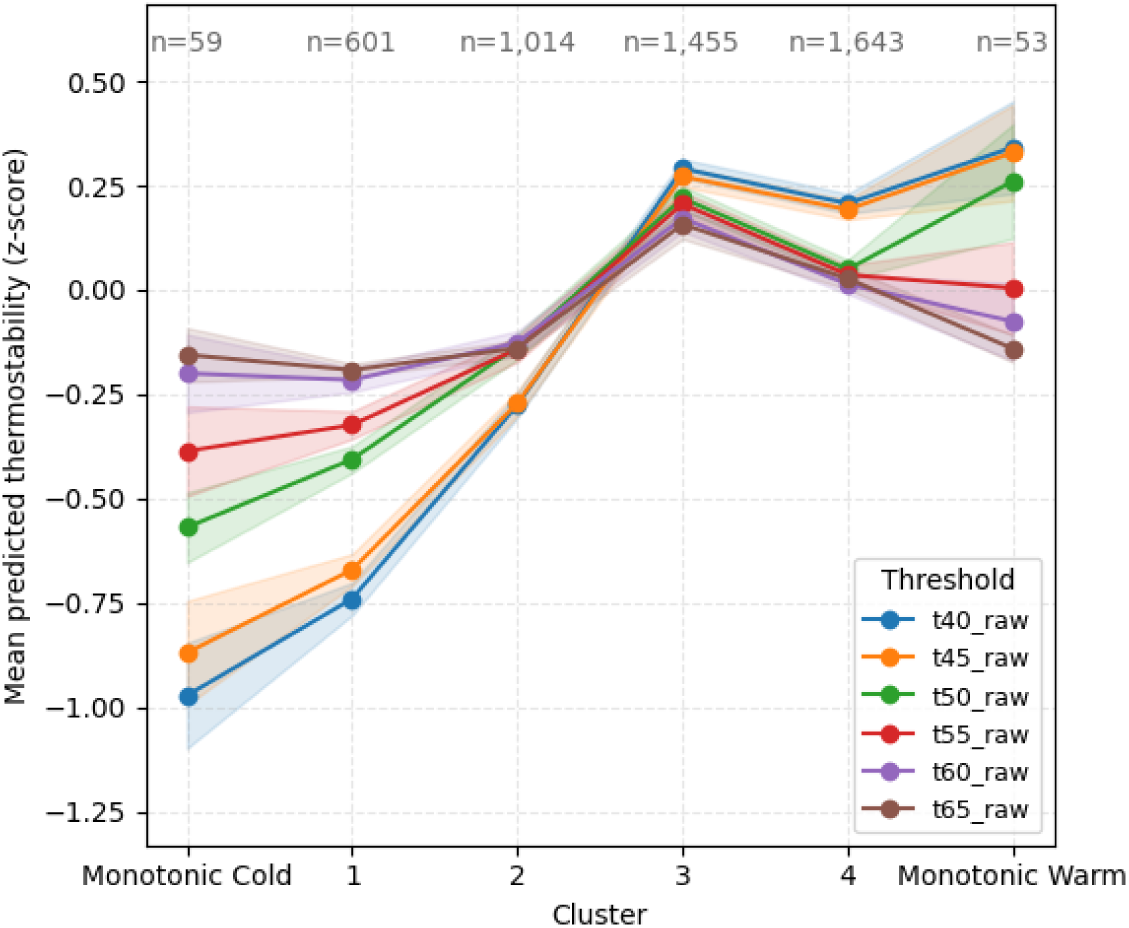
Predicted thermostability of COG0012 - Ribosome-binding ATPase YchF. Mean (z-scored) TemStaPro-predicted thermostability score at six temperature thresholds (t40-t65) for SCMG COG0012, plotted across the four thermal clusters and the two monotonic groups (Monotonic Cold, Monotonic Warm). Shaded bands show ±1 SEM. Sample sizes (n), i.e. number of mOTUs contributing background gene predictions, are shown above each group.

**Fig S20.**
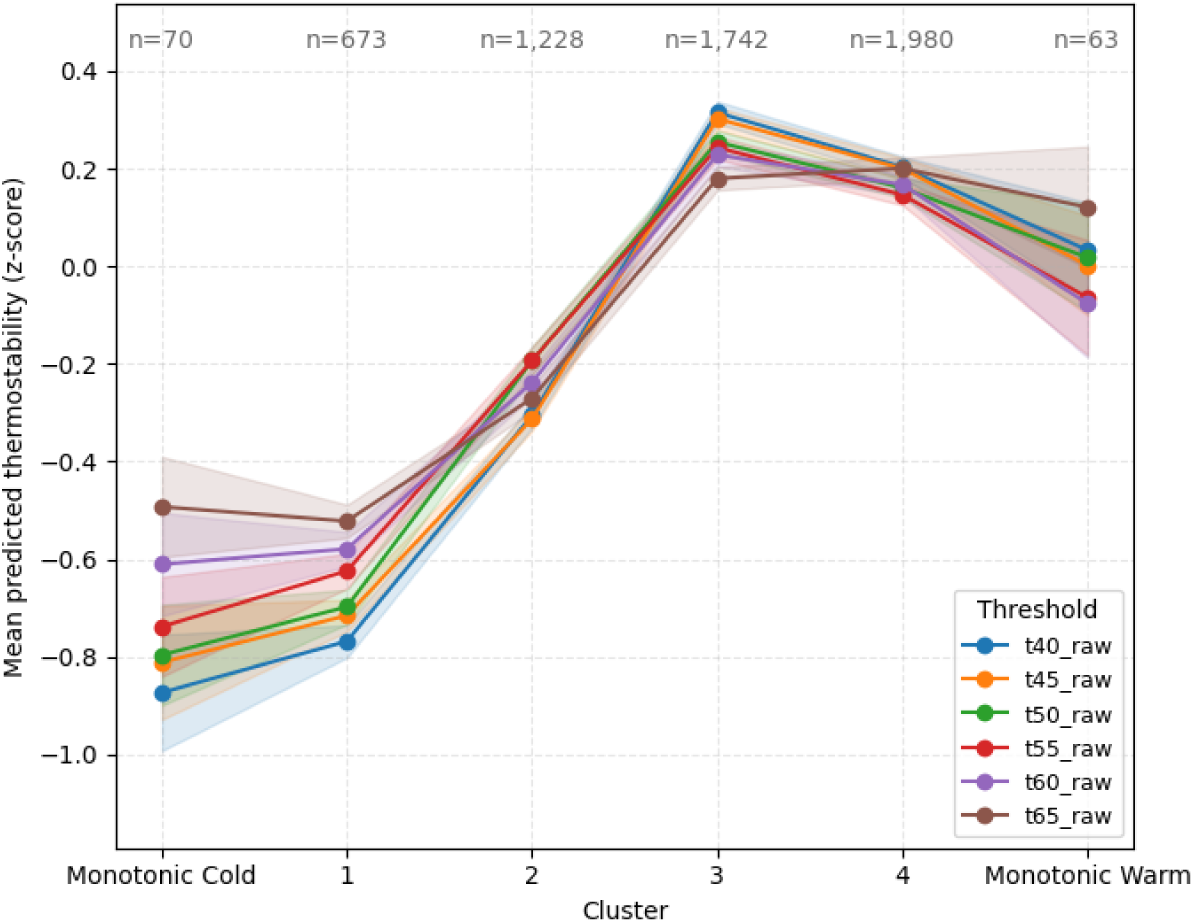
Predicted thermostability of a randomized background gene set across thermal clusters. Mean (z-scored) TemStaPro-predicted thermostability score at six temperature thresholds (t40-t65) for a background set of 100 randomly sampled protein sequences per mOTU representative genome (n = 5,756), plotted across the four thermal clusters and the two monotonic groups (Monotonic Cold, Monotonic Warm), ordered from coldest to warmest thermal association. Shaded bands show ±1 SEM. Sample sizes (n), i.e. number of mOTUs contributing background gene predictions, are shown above each group. This analysis parallels the SCMG-based prediction in Figure 4D, confirming that the cold-to-warm thermostability gradient is not restricted to single-copy marker genes but is detectable genome-wide.

**Fig S21.**
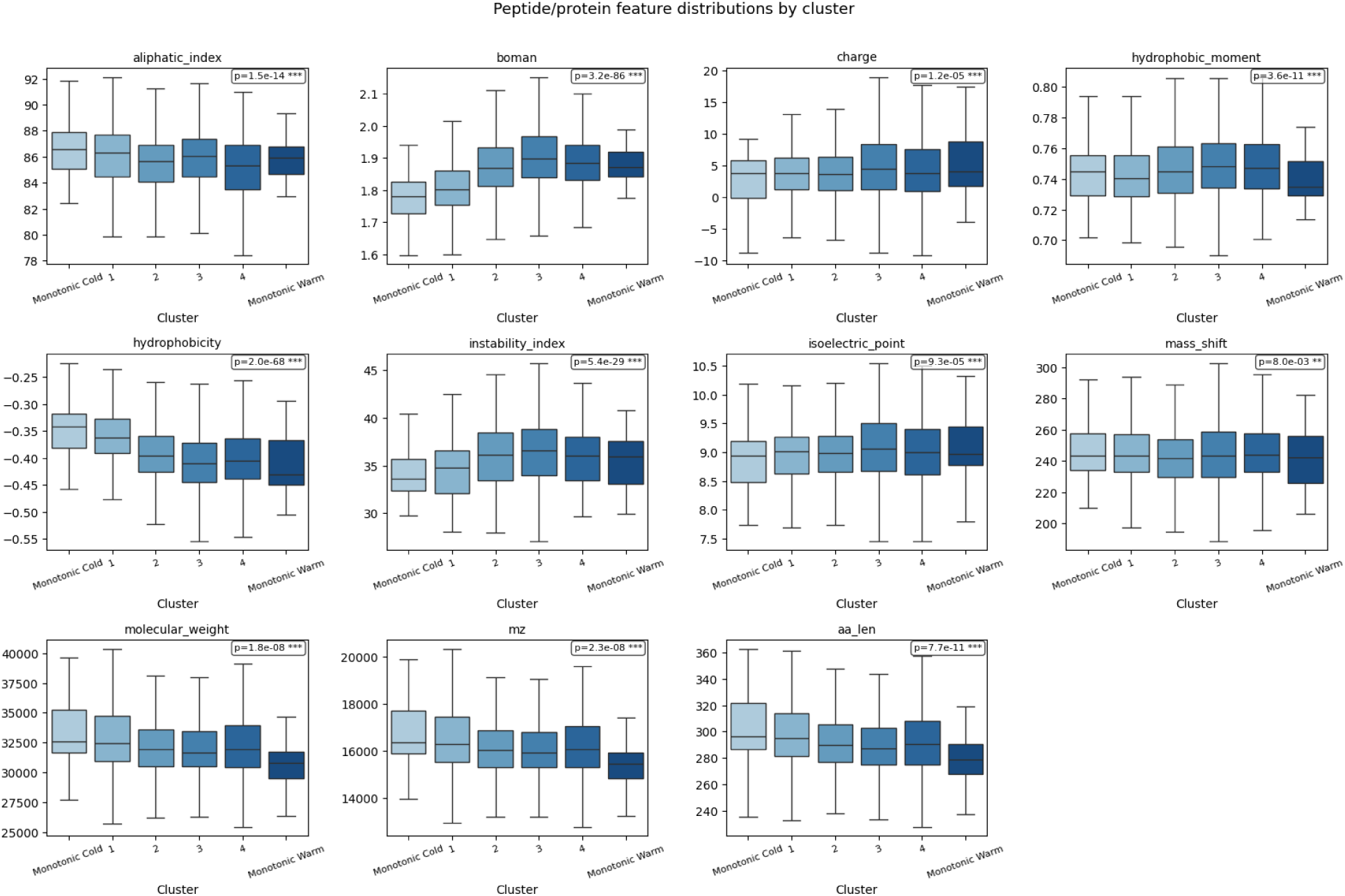
SCMGs physicochemical descriptors distributions by temperature trajectory cluster. Distribution of eleven physicochemical descriptors computed for SCMG-encoded proteins and compared across the four thermal clusters (ordered by ascending peak temperature). Boxes show the interquartile range and median; whiskers extend to 1.5× the interquartile range, with outliers omitted for clarity. p-values from Kruskal-Wallis tests are shown for each descriptor (*** p<0.001, ** p<0.01).

**Fig S22.**
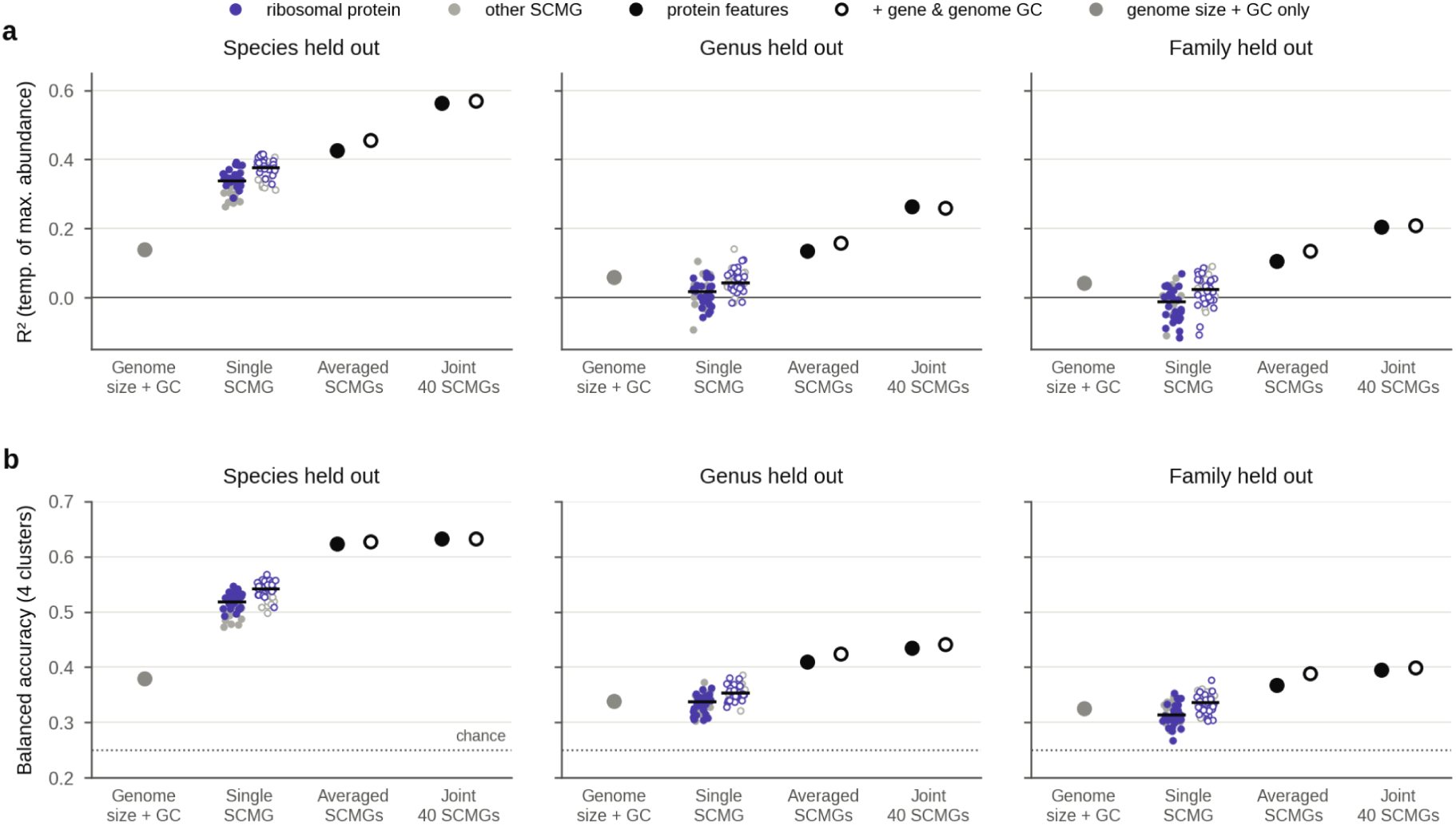
Predictive performance of protein feature sets across levels of phylogenetic independence. Out-of-fold performance of gradient-boosted models with mOTUs, genera or families held out of training (5-fold cross-validation; n = 5,756 mOTUs). (**A**) R^2^ for the temperature of maximum abundance; (**B**) balanced accuracy for the four thermal clusters (dotted line, chance). Models used genome size and GC content only, each SCMG individually (points; purple, ribosomal proteins; grey, other SCMGs; black bar, median of 40), the average of the 40 single-SCMG predictions, or all 40 SCMGs jointly. Filled symbols, eleven protein descriptors; open symbols, plus gene and genome GC content.

**Fig S23.**
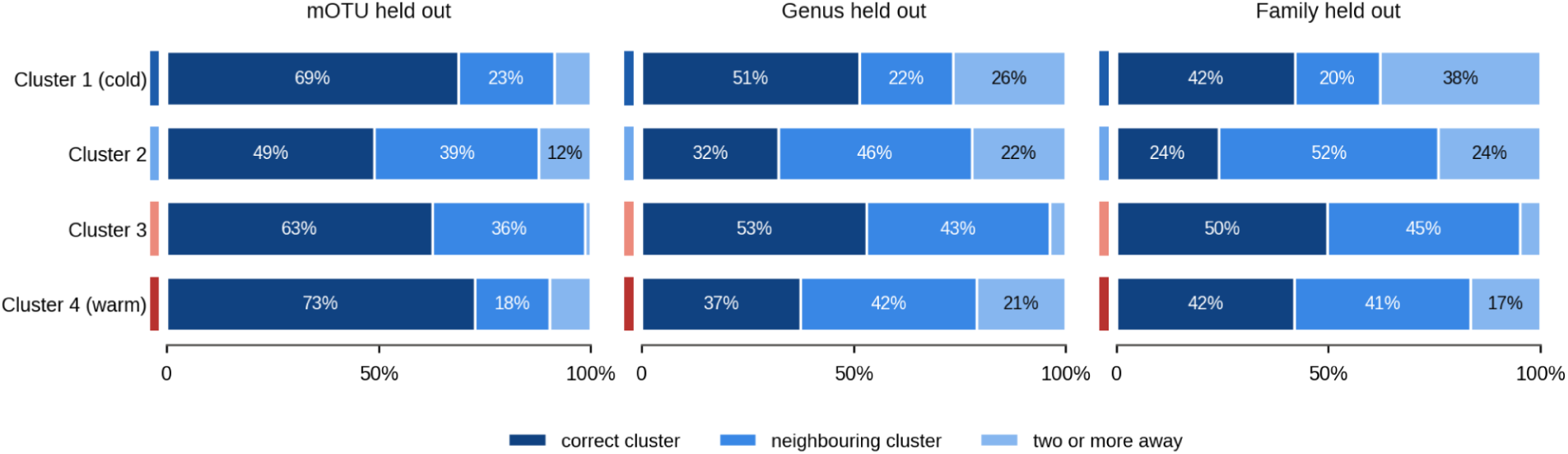
Classification of mOTUs into thermal clusters. Outcome of cluster assignment by the joint 40-SCMG classifier for each true cluster (correct cluster, neighbouring cluster, or two or more away), with mOTUs, genera or families held out of training.

**Fig S24.**
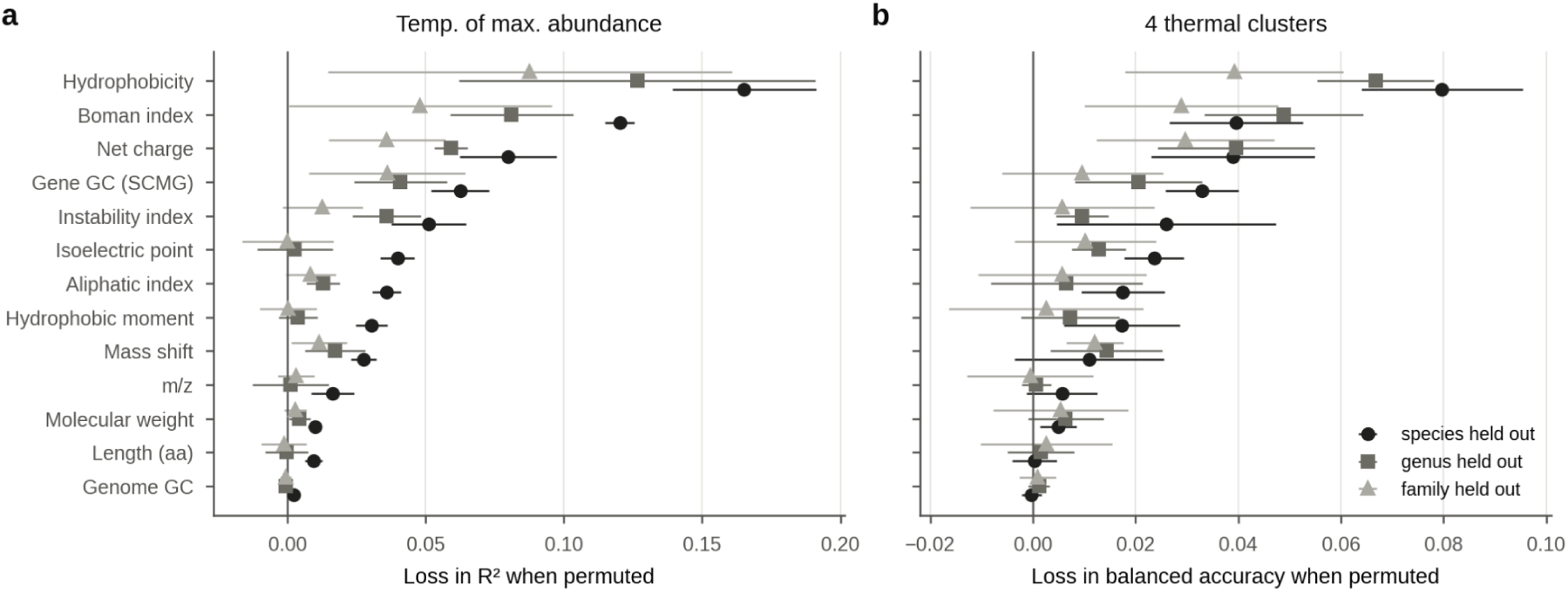
Contribution of each physicochemical descriptor to predictions. Grouped permutation importance of each descriptor in the joint 40-SCMG model (protein descriptors + GC). Each descriptor was permuted simultaneously across all SCMGs within each test fold. (**a**) Loss in R² for the temperature of maximum abundance; (**b**) loss in balanced accuracy for the four thermal clusters. Symbols show the mean ± SD across the five test folds, with mOTUs, genera or families held out.

## Notes

https://omdb.microbiomics.io/

### Competing Interest Statement

The authors have declared no competing interest.

